# Cryo-electron tomography sheds light on the largest protein in the *Trypanosoma brucei* tripartite attachment complex

**DOI:** 10.1101/2023.03.06.531305

**Authors:** Irina Bregy, Robin Benjamin Eberle, Julika Radecke, Amin Khosrozadeh, Jaime Llodrá, Akira Noga, Hugo van den Hoek, Mara Kern, Beat Haenni, Benjamin D. Engel, Philip Stettler, André Schneider, C. Alistair Siebert, Takashi Ishikawa, Benoît Zuber, Torsten Ochsenreiter

**Affiliations:** Institute of Cell Biology, University of Bern, Baltzerstrasse 4, Bern CH-3012, Switzerland; Institute of Anatomy, University of Bern, Baltzerstrasse 2, Bern CH-3012, Switzerland; Graduate School for Cellular and Biomedical Sciences, Freiestrasse 1, Bern CH-3012, Switzerland; Electron Bio-Imaging Centre, Diamond Light Source, Oxfordshire OX11 0DE, Didcot, United Kingdom; PSI Center for Life Sciences, Paul Scherrer Institute, Forschungsstrasse 111, Villigen CH-5232, Switzerland; Biozentrum, University of Basel, Spitalstrasse 41, Basel CH-4056, Switzerland; Department of Biology, ETH Zurich, Zurich CH-8092, Switzerland; Department of Chemistry Biochemistry and Pharmacy, Freiestrasse 3, Bern CH-3012, Switzerland

**Keywords:** Cryo-electron tomography (cryo-ET), tripartite attachment complex (TAC), mitochondrial genome anchoring, mitochondrial genome segregation, kDNA, *Trypanosoma brucei*

## Abstract

Trypanosomes only contain a single mitochondrion per cell. Within that singular mitochondrion, the protist carries a single mitochondrial genome that consists of a complex DNA network, the kinetoplast DNA (kDNA). The replicated kDNA is segregated during cell division by the tripartite attachment complex (TAC), a multi-protein bridge that physically links each daughter kDNA to a basal body (BB). BB movements drive kDNA segregation prior to cell division. How the TAC accommodates constant BB movements while maintaining a stable kDNA anchor throughout the cell cycle has remained unclear. Here we used cryo-electron tomography to image the cytoplasmic part of the TAC in its native context. We resolved the BB, the mitochondrial membranes, and the exclusion zone filaments (EZFs) connecting the BB and pro-BB to the outer mitochondrial membrane (OMM) and quantified the geometry of the region across many cells. EZF lengths spanned 230 to 625 nm in zoid cells and up to 874 nm in NP40-treated cells, and the BB occupied a wide range of positions and orientations relative to the OMM, while the pro-BB sat closer and more constrained. Building on these observations and prior evidence that p197 alone defines the BB-OMM distance, we propose that p197 is a length-variable connector: the length of a tandem array of α-helical repeats depends on their relative orientation, and filaments of different lengths coexist at one basal body. How that orientation is set remains open, but such a connector reconciles stable kDNA anchoring with the mechanical demands of BB movement during the cell cycle.

**Significance statement:** Using cryo-electron tomography, we describe structural features and dimensions of the tripartite attachment complex (TAC) that forms a stable link between the basal body and the mitochondrial DNA in *Trypanosoma brucei*. We show that p197, which spans this link and which we find to be the largest known TAC protein in *T. brucei*, forms filaments of different lengths at a single basal body, and we propose that it accommodates the forces of a moving basal body by varying its own length. The TAC is specific to trypanosomatids, but the problem of mechanical flexibility in dynamic cellular contexts is widespread and poorly understood. Previous work on mechanically compliant proteins has largely isolated molecules; cryo-electron tomography characterizes p197 *in situ*.

## Introduction

The mitochondrion is a hallmark of eukaryotic life, and it is responsible for essential processes ranging from catabolic reactions like oxidative phosphorylation to anabolic processes such as iron-sulfur cluster assembly (Braymer & Lill, 2017; Friedman & Nunnari, 2014). Many mitochondrial genes have been transferred to the nucleus, and thus the encoded proteins are produced in the cytoplasm and imported into the mitochondrion. Some essential genes, however, remain encoded on the mitochondrial genome. Consequently, trypanosomes, like most other eukaryotes, depend on the inheritance of an intact mitochondrial genome for their survival.

Along with the other members of the Kinetoplastea, trypanosomes possess one single, large mitochondrion per cell (Tyler, 2001). The singular nature of the mitochondrion and mitochondrial genome requires special care of replicating and segregating that genome correctly, prior to cytokinesis (Amodeo et al., 2023; Schneider & Ochsenreiter, 2018).

The mitochondrial DNA in trypanosomes is condensed to a disc-like network of interlocked circular DNA molecules that is positioned at the posterior end of the cell body (Jensen & Englund, 2012). This structure has been termed the kinetoplast, with its DNA content referred to as the kinetoplast DNA (kDNA) (Trager, 1965). The kDNA consists of two major classes of DNA molecules. The maxicircles (∼25 molecules of ∼23 kb) encode genes of the respiratory chain and the mitochondrial ribosome. Most of the genes are cryptogenes that require posttranscriptional RNA editing to encode functional proteins (Jensen & Englund, 2012; Ochsenreiter et al., 2007; Povelones, 2014). This editing process in turn required small noncoding gRNAs, which are encoded on the ∼5000 minicircles (∼1kb) (Cooper et al., 2019; Hong, 2003; Ochsenreiter et al., 2007). Within the kDNA network, minicircles are interlocked with each other and the maxicircles (Chen et al., 1995).

The main organizer of cell cycle progression in *T. brucei* is the basal body (BB), (Vaughan & Gull, 2015). To allow for coupling of kDNA segregation and BB separation prior to cytokinesis, the kDNA remains physically attached to the proximal end of the BB throughout the cell cycle (Ogbadoyi et al., 2003; Robinson & Gull, 1991; Schneider & Ochsenreiter, 2018). This connection, spanning the cytosol, the mitochondrial membranes and the mitochondrial matrix, has been termed as the tripartite attachment complex (TAC) (Ogbadoyi et al., 2003). The TAC not only allows for coordinated segregation of the kDNA but also anchors the kDNA to its position throughout the cell cycle. As the TAC has to bridge three biochemically distinct regions of the cell to reach from the BB to the mitochondrial DNA, it has been subdivided into three substructures: i) the exclusion zone filaments (EZFs) connecting the BB and the pro-basal body (pBB) to the outer mitochondrial membrane (OMM), ii) the differentiated mitochondrial membranes that show increased detergent resistance (DMs) and iii) the unilateral filaments (ULFs) that span the distance between the inner mitochondrial membrane (IMM) and the kDNA itself (Ogbadoyi et al., 2003).

Over the past decades, nine components of the TAC have been identified (Amodeo et al., 2023; Schneider & Ochsenreiter, 2018), with the most recent addition to this large protein complex being TAC53 (Jetishi et al., 2025). While most of them localize to the DMs or the ULFs, p197 spans the entire exclusion zone (Aeschlimann et al., 2022; Gheiratmand et al., 2013). Its N-terminus localizes at the OMM, while the C-terminus localizes to the proximal end of the (p)BB (Aeschlimann et al., 2022). In addition, p197 contains a central section of more than 25 nearly identical repeats of 175-182 amino acids (Aeschlimann et al., 2022; Naguleswaran et al., 2021). p197 stably anchors the BB to the DMs while accommodating the extensive remodeling of this region during the *T. brucei* cell cycle, and the extent of the repeat section suggests a central role in this mechanical function. Interestingly, the amino acid sequence of the ortholog of p197 found in the closely related parasite *Trypanosoma cruzi* (Tcp197) is more than 70% similar in the terminal domains but does not contain a repeat section in the central α-helical part of the protein (Aeschlimann et al., 2022). Consequently, the *T. cruzi* protein is much shorter and replacement of the *T. brucei* p197 repeat section with the corresponding central region of the *T. cruzi* ortholog demonstrates that the *T. cruzi* protein cannot complement for the function of the repeat section in *T. brucei* p197 (Aeschlimann et al., 2022). This difference in the composition of p197 is also reflected by differences in cell cycle dynamics observed between the two parasites. While the cell cycle of *T. brucei* has been shown to include a rotation of the new BB around the old BB prior to kDNA segregation, a similar process seems not to take place in *T. cruzi* (Elias et al., 2007; Vaughan & Gull, 2015). In fact, *T. cruzi* delays migration and repositioning of the new BB until asymmetric cell division is complete (Elias et al., 2007).

To better understand how p197 functions, we investigated the structural organization of the exclusion zone *in situ* using cryo-electron tomography (cryo-ET).

## Results

To gain an overview of the exclusion zone filaments (EZFs) of the TAC *in situ*, we focused on the procyclic form (PCF) of *T. brucei*, which corresponds to the insect stage of the parasite and is the established model for studies of TAC biology. We used cryo-ET on vitrified PCF trypanosomes. Since *T. brucei* cells are above 500 nm in thickness in the region of the TAC, we used several different approaches to improve the contrast in that region. The most successful approaches were (i) the gentle treatment with detergent (NP40) (Figure 1, Figure S1) and (ii) the generation of “zoid” cells (trypanosomes that have lost their nucleus following depletion of TbCentrin4 (Aeschlimann et al., 2022; Amodeo et al., 2023; Cooper et al., 2019; Elias et al., 2007; Gheiratmand et al., 2013; Hajduk & Ochsenreiter, 2010; Hong, 2003; Jensen & Englund, 2012; Jetishi et al., 2025; Naguleswaran et al., 2021; Ochsenreiter et al., 2007; Ogbadoyi et al., 2003; Robinson & Gull, 1991; Schneider & Ochsenreiter, 2018; Shi et al., 2008; Sun et al., 2018; Trager, 1965; Vaughan & Gull, 2015), which are thin enough for cryo-ET (Figure 2 and Figure S2). We also visualized the EZFs in untreated whole cells and in cryo-focused ion beam milled (cryo-FIB) cells. However, in both cases, the number of TAC regions that could be identified and visualized was very low (Figure S2).

**Figure 1:**
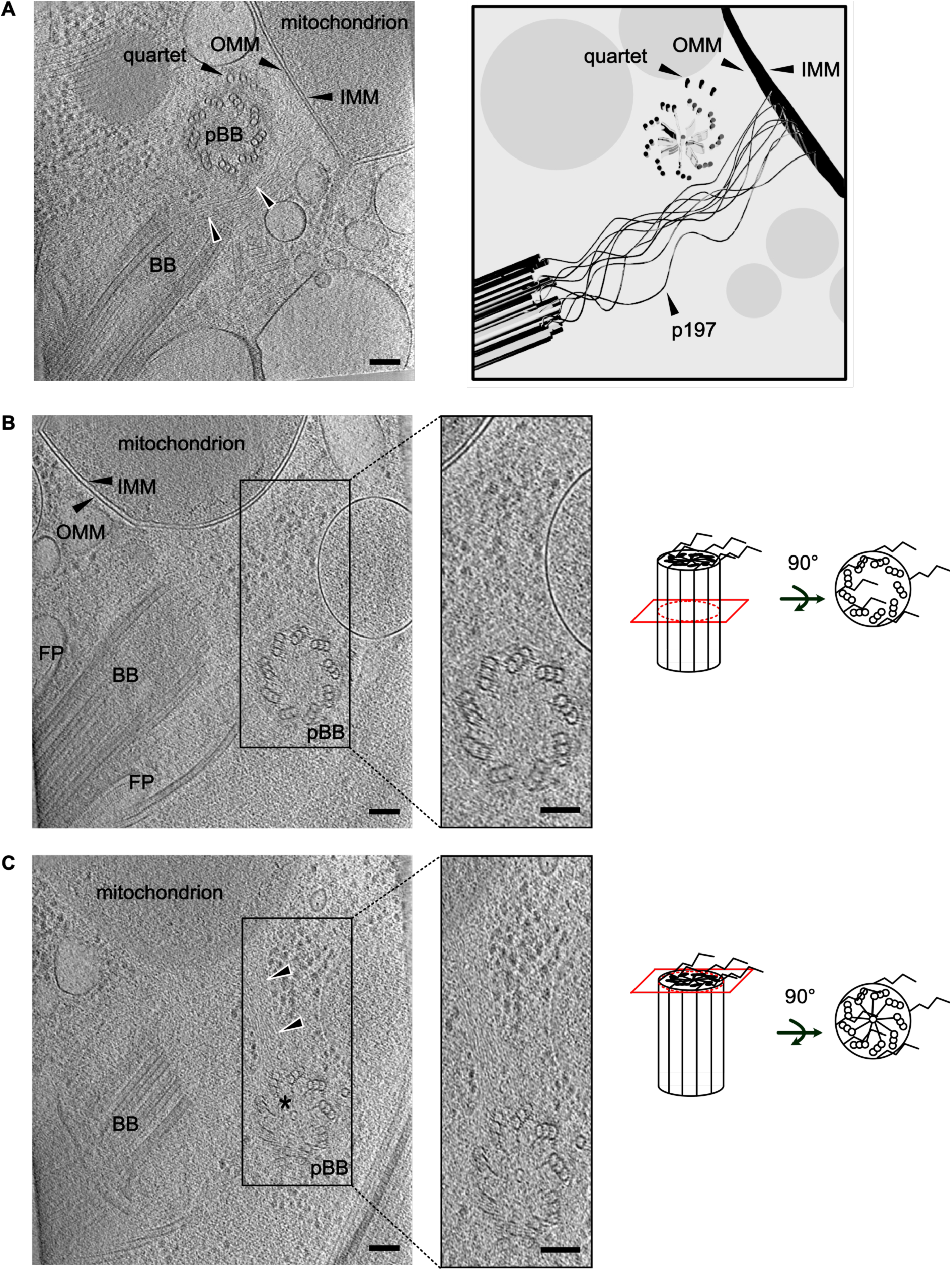
Architecture of the TAC region visualized by NP40 treated whole-cells cryo-ET of detergent-treated *T. brucei*. Cells were briefly treated with 0.5% NP40 prior to plunge freezing, which preserves the ultrastructure of the mitochondrial membranes and cytosolic ribosomes while improving contrast in the TAC region. A) z-slices (n = 10) through a representative tomogram of the TAC region, showing the basal body (BB), pro-basal body (pBB), p197 filaments (black arrowheads with white border), the outer and inner mitochondrial membrane (OMM, IMM), the flagellar pocket (FP), and the emerging microtubule quartet (visible is a triplet only). Right: Schematic 3D rendering of the same region. For visualization purposes, the model is rotated and therefore does not exactly correspond to the view in the tomogram. B) z-slices (n = 10) through a different cell, showing a cross-section through the middle of the pBB and a longitudinal section of the BB and the base of the axoneme within the flagellar pocket (FP). Zoom: magnified view of the marked area, showing ribosomes but no p197 filaments between the pBB and the OMM. *Schematic on the right*: the red rectangle indicates the position of the z-slice through the pBB cylinder; rotation by 90° reveals the cross-section view as it appears in the tomogram. Black filaments on top represent p197 emerging from the proximal end of the pBB. C) Another z-slice (n = 10) through the same cell as in (B) cross-sectioning the proximal end of the pBB as indicated by the characteristic cartwheel structure with SAS-6 (asterisk). p197 filaments extend from the pBB toward the OMM, locally excluding ribosomes. Bars, 100 nm

**Figure 2:**
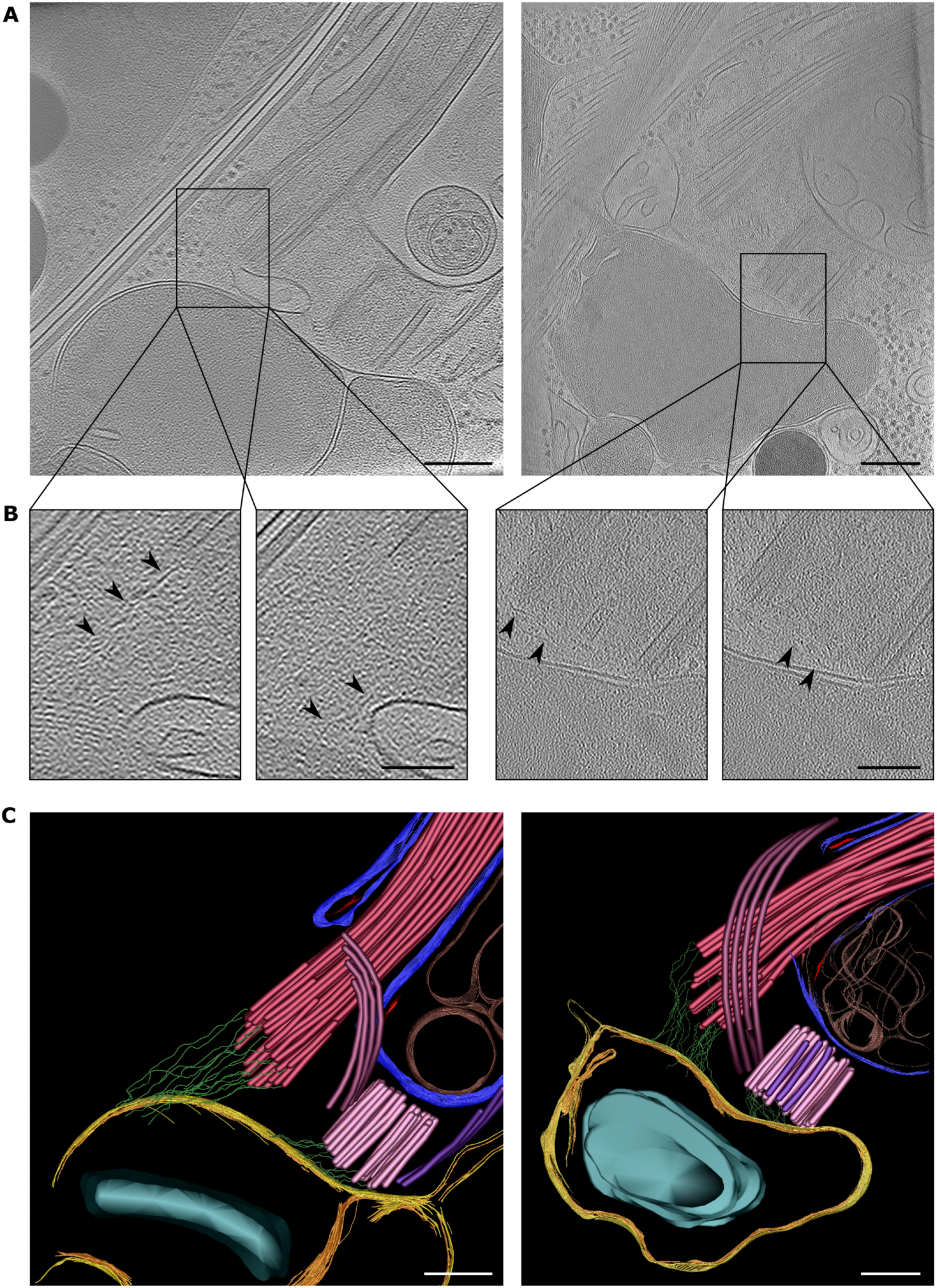
Cryo-ET of the zoid TAC area of two selected tomograms. A) Two representative cryo-electron tomograms of the TAC region, each shown as a single z-slice (right tomogram: 20.6 nm z-slice). Both scale bars: 200 nm. B) Zoom-in of the exclusion zone of the tomograms shown above, highlighting individual EZFs. Arrowheads point at the most well-resolved filament in the respective image. Scale bars: 100 nm. C) Model of the TAC area based on segmentations of the tomograms shown above. BB and flagellum (magenta), pBB (pink), microtubule quartet of the mature BB (purple), growing microtubule quartet of the pBB (violet), flagellar pocket (blue), flagellar pocket vesicles (brown), collarette (red), exclusion zone filaments of the TAC (green), outer mitochondrial membrane (yellow), inner mitochondrial membrane (orange) and kDNA (cyan) have been segmented in cryo-CARE (Buchholz et al., 2018) denoised tomograms. The flagellar pocket is a membranous invagination through which the flagellum exits the cell body; the microtubule quartet is a set of four parallel microtubules that emerge near the basal body wrap around the flagellar pocket. Scale bars: 200 nm.

The brief treatment of PCF *T. brucei* cells with 0.5% NP40-containing buffer prior to plunge freezing and cryo-ET preserved the ultrastructure of the mitochondrial membranes, cytosolic ribosomes and the microtubules with their protofilaments, while yielding high contrast, as can be seen in the cross section of the pro-basal body (Figure 1A). We detected the inner and outer mitochondrial membranes (IMM and OMM), the basal body (BB) and its associated flagellar pocket, and the pro-basal body (pBB) with its emerging microtubule quartet (Figure 1A). We observed filaments between the proximal end of the BB and the OMM, which likely correspond to the protein p197 (Figure 1A). A schematic representation of this region is shown in the right part of Figure 1B. Although filaments were consistently visible in all tomograms containing the TAC region, their small diameter and their high local density, combined with the limited signal-to-noise ratio of cryo-electron tomograms, prevented automatic tracing and required manual segmentation. Figure 1B and C display different sections of a single tomogram. In Figure 1B, both mitochondrial membranes are visible along with a longitudinal section through the BB and a cross-section of the central region of the pBB, which is positioned at a 90° angle relative to the BB. Between the pBB and the OMM, only cytosolic ribosomes are visible; no filaments are detected. Moving in the tomogram towards sections that contain the proximal end of the pBB reveals the characteristic cartwheel structure, with the SAS6 hub at its center (Figure 1C) (Kitagawa et al., 2011; van Breugel et al., 2011). From this region of the pBB, filaments extend towards the OMM. The filaments exclude ribosomes from the area, giving rise to their original designation as exclusion zone filaments (Ogbadoyi et al., 2003). We also detected the filaments in Triton X-100-extracted cells. This stronger treatment extracts most cytoplasmic content and solubilizes most membranes, leaving clearly visible only the basal body, parts of the axoneme, the pro-basal body, and the filaments emanating from the proximal end of the basal body (Figure S2E).

In summary, brief detergent treatment yields high-contrast tomograms of the TAC region. The exclusion zone filaments are not only associated with the mature basal body but are also present at the pro-basal body, where they extend towards the OMM and locally exclude cytosolic ribosomes (Figure 1B, C and Figure S1).

While the TAC area was well resolved in the detergent-treated cells, we also aimed to visualize the structure under native conditions. In untreated whole cells and cryo-FIB-milled cells, we visualized the TAC in only a few cases (Figure S2A, F). We therefore turned to zoid cells, whose reduced thickness enables cryo-ET without the need for FIB-milling.

### Morphology of the TAC area in zoids

We prepared the zoids as described previously and verified the presence of the TAC using immunofluorescence and transmission electron microscopy see figure S2D, S3, S4 (Shi et al., 2008). In most zoid tomograms, we readily identified the TAC and surrounding structures such as the pBB, the BB, the flagellar pocket, the microtubule quartet, and the mitochondrial kDNA pocket (Figure 2).

As in the detergent-treated cells (Figure 1), we identified the exclusion zone filaments between the proximal ends of the (p)BB, in a region free of ribosomes (arrowheads in Figure 2B, 3D-segmentation in Figure 2C). The filaments are thin and wavy, and span the entire region between the BB and the OMM, as well as between the pBB and the OMM. To confirm that the filaments were indeed part of the TAC, we acquired over 30 tomograms of tdzoid cells that had also been depleted of the p197 protein by RNAi (tdzoids, Figure S3D). We found no filamentous structures between the (p)BB and the OMM in any of these tomograms; instead, the area contained ribosomes, like the rest of the cytoplasm (Figure S5). In the absence of p197, the exclusion zone and its filaments are abolished.

**Figure 3:**
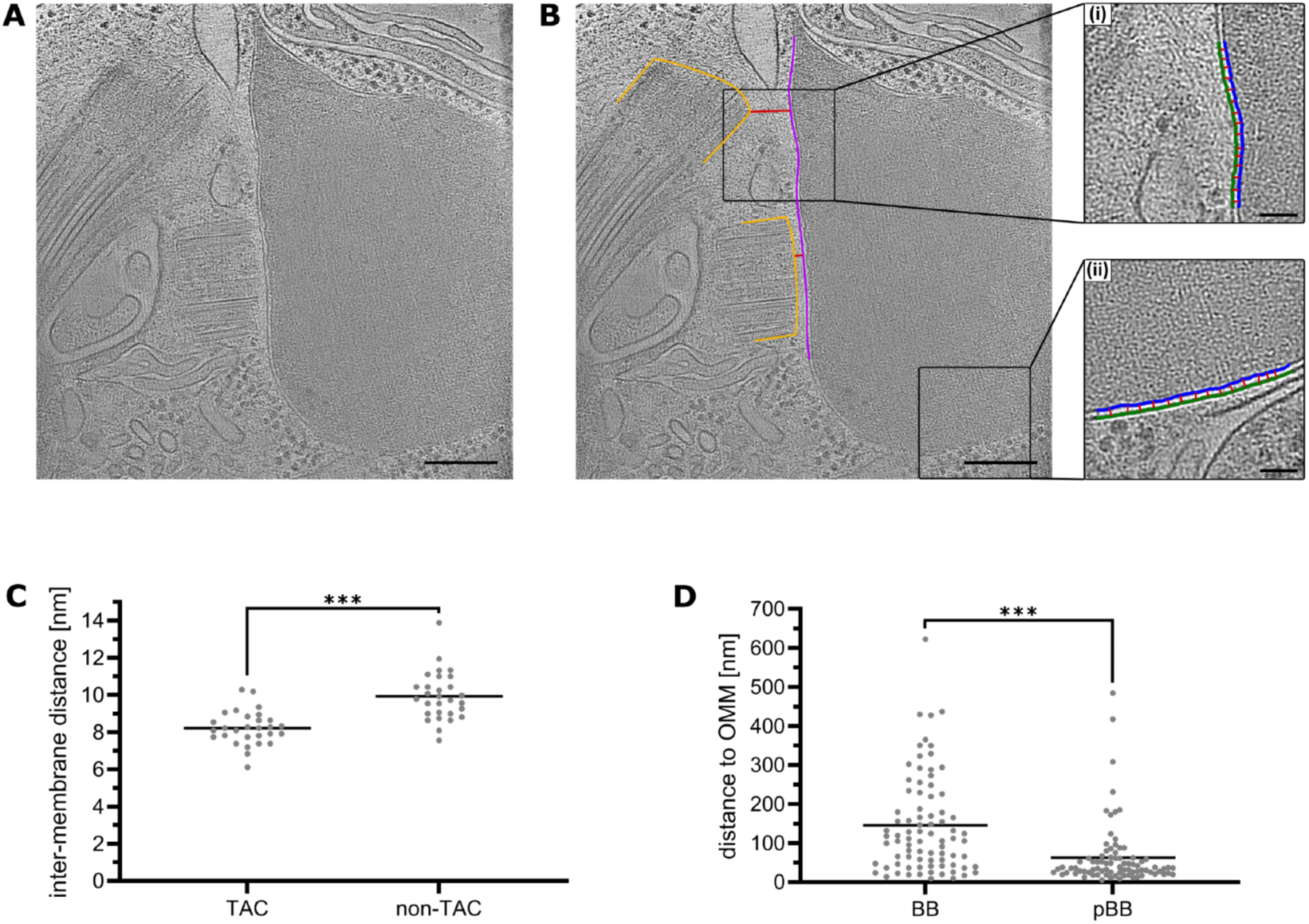
Geometry of the TAC region: inter-membrane distances and basal body positioning relative to the outer mitochondrial membrane. A) Single z-slice (1.03 nm) of a representative tomogram of the TAC area in a *T. brucei* zoid cell. Scale bar: 200 nm. B) Same z-slice as in (A) with annotations illustrating the measurement strategy. The orange contour outlines the proximal face of the BB and pBB; the magenta line follows the OMM. Red lines indicate the measured BB-OMM and pBB-OMM distances (quantified in (D)). The two boxed regions show membrane patches selected for intermembrane distance measurements: one within the TAC area (inset i, top, from the same z-slice) and one outside the TAC area (inset ii, bottom, from a different z-slice of the same tomogram). In the insets, the OMM is traced in blue, the IMM in green, and red lines indicate the local intermembrane distance (measured between the centers of the two membranes). Scale bars: 200 nm (B), 50 nm (insets). C) Inter-membrane distance in the TAC area compared to the area outside of the TAC (non-TAC). Each point represents the average of 15 measurements per membrane patch in one cell (n = 29 cells). Horizontal bars indicate the mean. *** p ≤ 0.001 (two-sided paired t-test, p = 5 × 10⁻¹⁰). D) Distance to the OMM measured for the BB and the pBB (n = 82 cells). Horizontal bars indicate the mean. *** p ≤ 0.001 (two-sided paired t-test, p = 1.4 × 10⁻⁶).

Inside the mitochondrion, we did not observe any filamentous structures corresponding to the ULFs, i.e. intramitochondrial components of the TAC. However, the intermembrane distance, measured at 15 positions per cell in 29 cells, was significantly smaller in the TAC area (8.2 ± 0.9 nm) than outside of it (9.9 ± 1.3 nm) (Figure 3A-C).

We measured the minimal distance between the OMM and the proximal face of either the BB or the pBB (Figure 3D, n = 82). pBBs were located on average 63 ± 81 nm away from the OMM, significantly closer than the mature BB (145 ± 123 nm). In 70% of the cells, the pBB was closer to the OMM than the mature BB, and the spread of the distance data was also larger for the BB. This difference in spread was confirmed statistically (Levene’s test, p = 7 × 10⁻⁵; variance ratio = 2.34; Brown-Forsythe test gave an identical result).

Collectively, these data confirm that a p197-dependent filamentous structure connects the (p)BB to the OMM in zoid cells. They also reveal a broad range of BB and pBB positions relative to the OMM.

### Basal body and pro-basal body orient independently relative to the outer mitochondrial membrane

In addition to the variable BB and pBB positions relative to the OMM described above, we noticed that the orientation of both axes relative to the OMM was also highly variable (Figure S6). To quantify this variability and to detect any preferred orientations, we measured the inclination of each axis relative to the OMM plane, as well as the angle between the BB and pBB axes (Figure 4A). We defined 0° as an axis lying in the OMM plane and 90° as an axis perpendicular to it. Both axes were on average steeply inclined toward the OMM, with mean inclinations of 48.8 ± 23.6° for the BB and 51.1 ± 25.1° for the pBB (n = 73; Figure 4B, C). These two distributions were statistically indistinguishable (Wilcoxon signed-rank test, p = 0.228), but they spanned almost the full possible range, from nearly parallel to the membrane to nearly perpendicular. The inter-axis BB-pBB angle showed a comparably widespread (mean 51.4 ± 39.8°, range from close to 0° to above 150°), but with a distinctly asymmetric distribution: small angles (BB and pBB nearly aligned) were the most frequent configuration, while large angles (BB and pBB splayed apart or nearly anti-parallel) were progressively rarer (Figure 4D).

**Figure 4:**
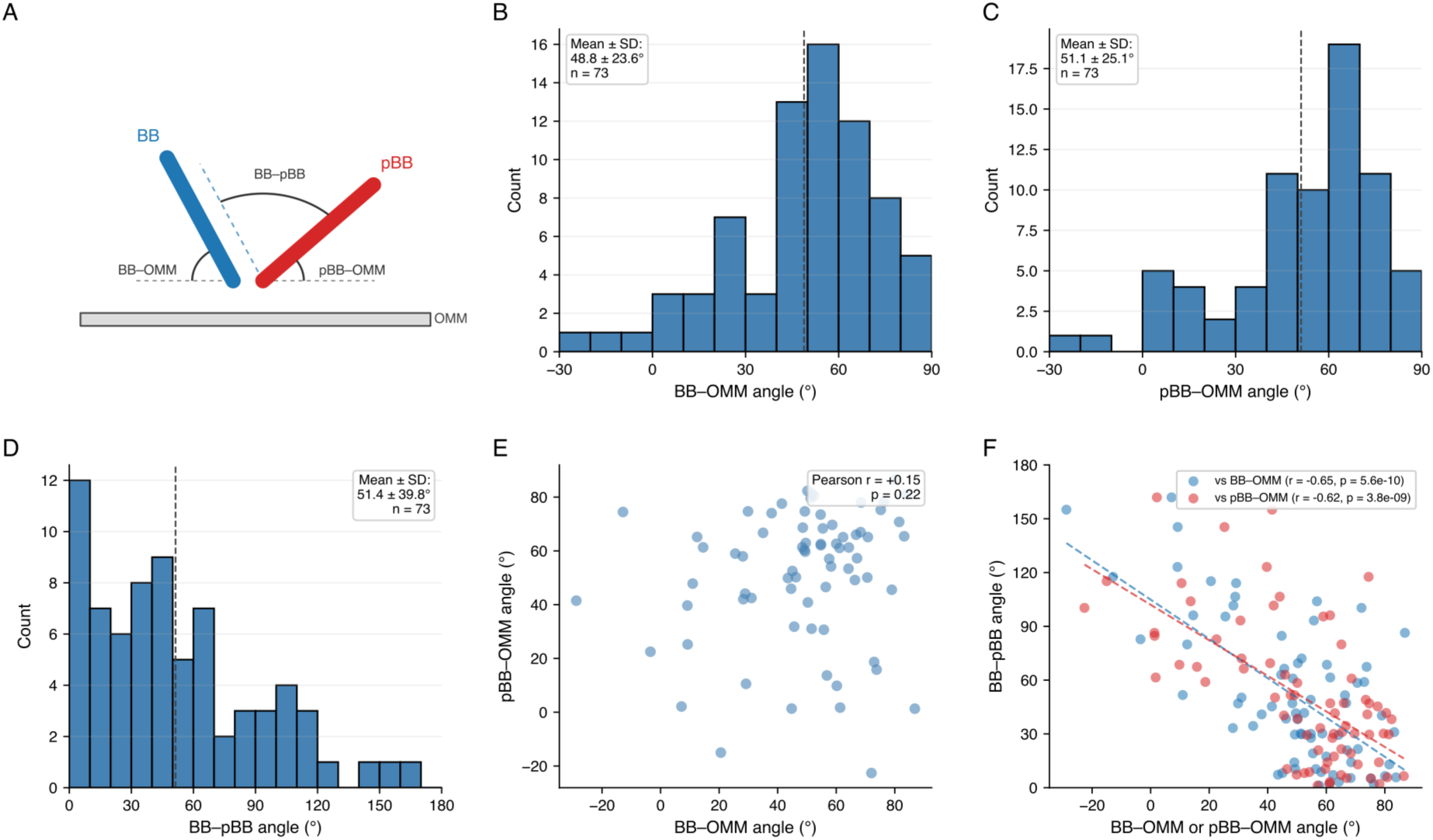
Orientation of the basal body and pro-basal body relative to the outer mitochondrial membrane. A) Schematic definition of the three angles measured. The BB-OMM and pBB-OMM angles are the inclinations of the BB and pBB axes relative to the OMM plane in the TAC region (0° = axis lying in the membrane plane; +90° = axis perpendicular to the membrane and pointing toward the cytoplasm). The BB-pBB angle is the unsigned angle between the two axes (0° = parallel; 180° = anti-parallel). See Methods for the geometric definitions and sign convention. B-D) Distribution of the three angles across 73 tomograms. Mean ± SD is indicated in each panel; the dashed vertical line marks the mean. B) BB-OMM angle. C) pBB-OMM angle. D) BB-pBB angle. E) BB-OMM angle plotted against pBB-OMM angle for each tomogram. Pearson correlation coefficient and p-value are indicated. F) BB-pBB angle plotted against the BB-OMM angle (blue) and the pBB-OMM angle (red), with linear regression lines. Pearson correlation coefficients and p-values are indicated for each series.

Despite both axes sampling a similar range of orientations, their inclinations toward the OMM were uncorrelated (Pearson r = 0.15, p = 0.22, Figure 4E): knowing how steeply one axis is tilted toward the membrane provides no significant linear correlation between the two inclinations. In contrast, the inter-axis BB-pBB angle was strongly and negatively correlated with both membrane-inclination angles, and to a similar extent (BB-pBB vs BB-OMM: Pearson r = -0.65, p = 5.6 × 10⁻¹⁰; BB-pBB vs pBB-OMM: Pearson r = -0.62, p = 3.8 × 10⁻⁹; Figure 4F). In other words, when either axis was steeply inclined toward the membrane, the BB and pBB tended to be closely aligned with each other; when either axis lay more parallel to the OMM, the two axes splayed further apart. The near-identical slopes of the two correlations (Figure 4F) indicate that this geometric coupling is symmetric: it operates equivalently through the BB and the pBB.

Taken together with the wide range of BB-OMM and pBB-OMM distances reported above, these data show that the BB and pBB can adopt a wide range of positions and orientations relative to the OMM, despite being stably anchored to the mitochondrial membrane through the TAC. This variability is consistent with a flexible, possibly elastic nature of the exclusion zone filaments, which would allow them to maintain the anchorage while accommodating the observed range of configurations.

### Structural analysis of individual exclusion zone filaments

Having shown that p197 is required for EZF formation (Figure S5), we next examined how its molecular organization could give rise to the observed filamentous architecture. Previous work established that p197 localizes to the TAC and is essential for TAC formation and kDNA segregation (Gheiratmand et al., 2013; Hoffmann et al., 2018). Ultrastructure expansion microscopy revealed that p197 spans the entire exclusion zone, with its N-terminus at the DMs and its C-terminus at the BB and pBB (Aeschlimann et al., 2022). Using Southern blot analysis on a p197 allele harboring a silent recoded region for specific probe detection, we found that the p197 locus contains 34.5 repeats (68% CI: 31.5 – 38.0; Figure S7), substantially more than the ∼3.5 reported in the original genome assembly (Berriman et al., 2005) or the ∼26 inferred from a long-read gDNA assembly (Naguleswaran et al., 2021). Given the size of one repeat (175-182 amino acids), this corresponds to an estimated gene size of ∼21.7 kb and a predicted protein mass of ∼796 kDa, making p197 the largest known TAC protein in *T. brucei*.

In 22 tomograms from zoid cells, we manually segmented and measured five traceable EZFs each, giving a total of 110 length measurements (Figure 5A, B). Because filaments traced within one tomogram are not independent, we treated the tomogram as the unit of analysis throughout. Mean EZF length was 417.5 ± 68.5 nm (mean ± SD of 22 tomogram means; 95% CI: 387 to 448 nm), while individual filaments spanned 229.6 to 624.5 nm. Within individual tomograms, the five filaments differed by 158 nm on average and by up to 282 nm, with the longest filament being on average 1.5 times the shortest, a spread as large as that between cells. Since all five filaments were traced in one vitrified cell, this spread cannot reflect differences between cells or between cell cycle stages: p197-dependent filaments with different traced contour lengths coexist at a single basal body. In 12 tomograms of NP40-treated wild-type cells, in which every resolvable filament was traced (430 in total), mean EZF length was 463.6 ± 144.5 nm (95% CI: 372 to 555 nm), with individual filaments spanning 29.3 to 874.3 nm. No minimum length was imposed for the dataset of the NP40 treated cells; the lower end of the NP40 range reflects that every resolvable filament, including short or partial contours, was traced, whereas the zoid dataset comprised only the five most clearly resolved filaments per tomogram. Compared on a per-tomogram basis, the two preparations gave indistinguishable mean filament lengths (Welch’s t test, p = 0.32; Mann-Whitney U test, p = 0.63), while the NP40 dataset was more dispersed (Levene’s test, p = 0.005), as expected from tracing every resolvable filament rather than the five best-resolved ones per tomogram (Table 1).

**Figure 5:**
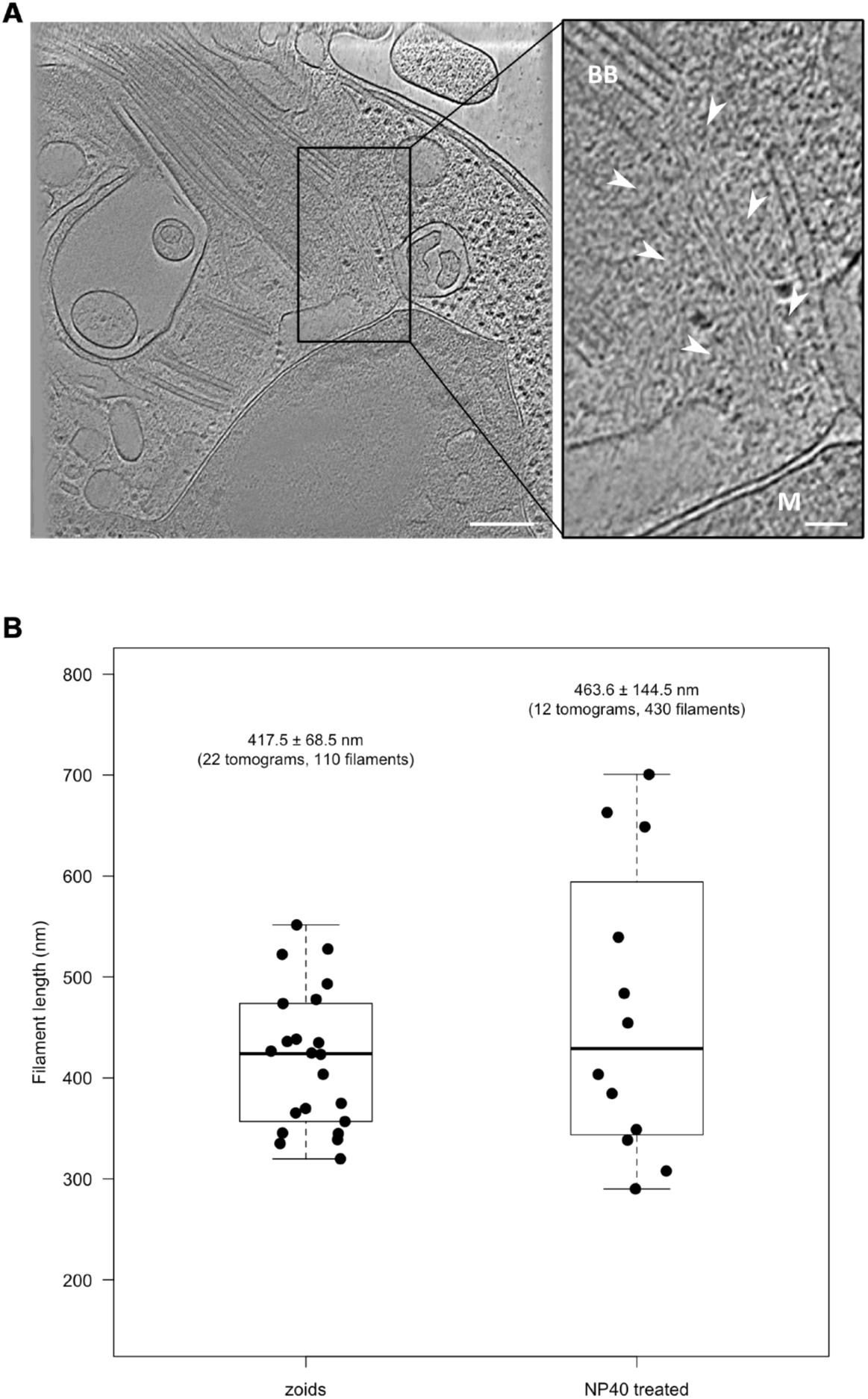
Length distribution of p197 filaments. A) Representative tomogram showing p197 filaments in zoid cells. Left: An overview of the TAC area with the boxed region shown enlarged on the right. Right: Enlarged view with white arrowheads surrounding the region of well resolved p197 filaments. M, mitochondrion; BB, basal body. Scale bars: 200 nm (overview), 50 nm (enlargement). B) Comparison of p197 filament lengths measured under two preparations. zoids (left; n = 110 filaments from 22 tomograms, 5 filaments manually traced per tomogram) and NP40-treated whole cells (right; n = 430 filaments from 12 tomograms, comprehensively traced per tomogram). Each point represents the mean filament length within one tomogram. Boxes show the interquartile range of tomogram means; the horizontal line within each box indicates the median. Labels give the pooled mean ± SD and total filament count for each dataset.

**Table 1.** Summary of p197 filament length measurements from two independent tomogram datasets. Zoids’ dataset consisted of 5 manually traced filaments per tomogram across 22 tomograms (110 filaments total); The dataset from the NP-40 treated cells consisted of comprehensively traced filaments (11–88 per tomogram) across 12 tomograms (430 filaments total).

|  |  |  |
| --- | --- | --- |
| Preparation | zoids | NP40 treated |
| Tomograms | 22 | 12 |
| Filaments | 110 (5 per tomogram) | 430 (11 to 88 per tomogram) |
| Mean ± SD of tomogram means | 417.5 ± 68.5 nm | 463.6 ± 144.5 nm |
| 95% CI of the mean | 387 to 448 nm | 372 to 555 nm |
| Individual filament range | 229.6 to 624.5 nm | 29.3 to 874.3 nm |

Since no experimental structure of p197 is available, we examined structure predictions of the repeat region. The AlphaFold Database entry for p197 (Jumper et al., 2021; Varadi et al., 2024) models the N- and C-terminal regions as unstructured and the repeat region as a series of α-helical stretches. To test this at higher resolution, we modeled the 175-residue repeat unit alone and in tandem arrays of up to six repeats using AlphaFold 2 and AlphaFold 3 (Figure 6, Figure S8, Table S1, Table S2). Modeled on its own, the repeat is predicted as one continuous α-helix in all 30 models of Figure 6A, from both programs and two registers of the repeat period, at a mean pLDDT of 83.3 to 90.3. A third register, cut in the middle of the helix, gives the same continuous helix in 14 of 15 models, though at lower confidence and, in AlphaFold 2, far more compact (Table S1).

**Figure 6:**
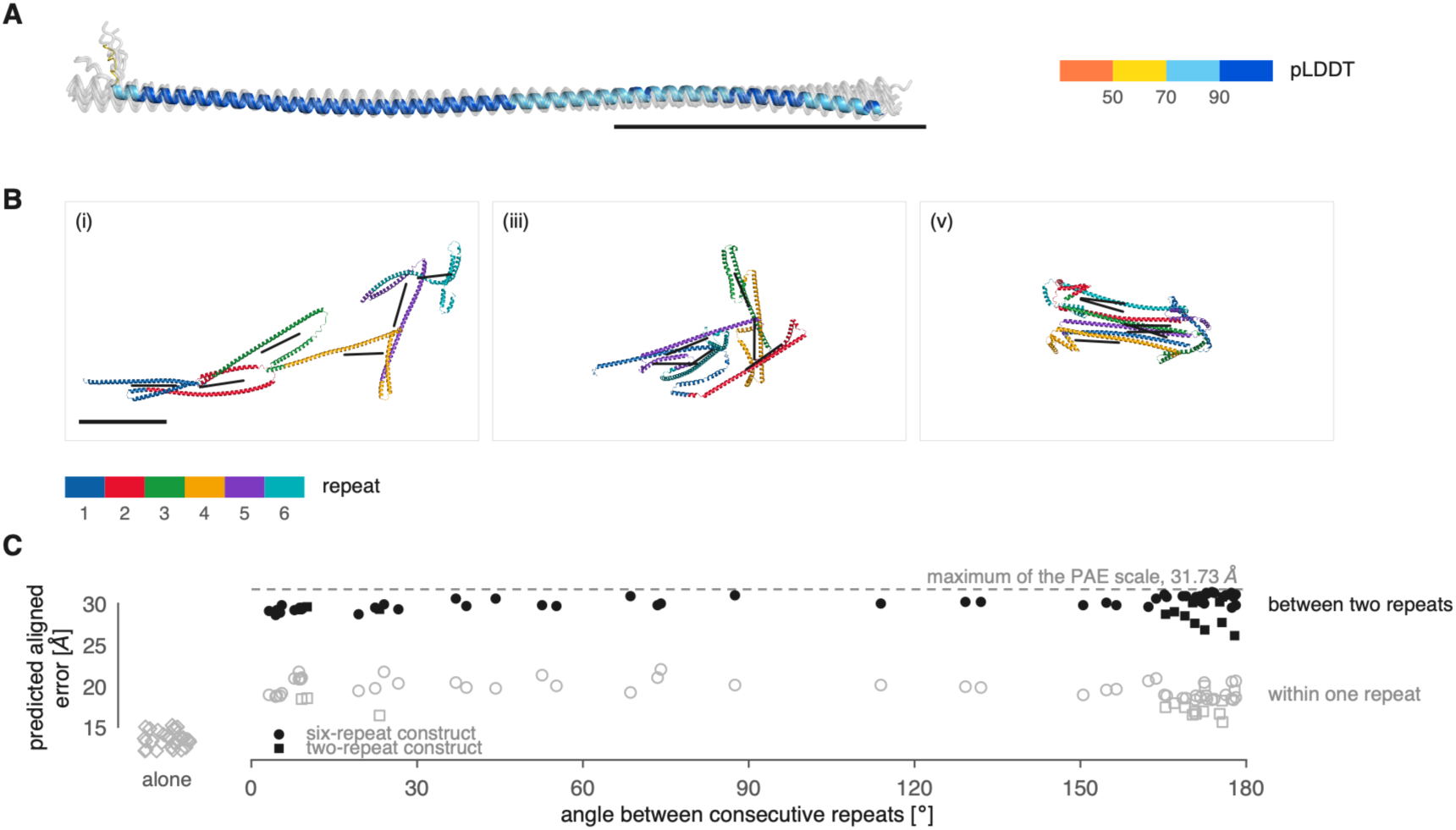
AlphaFold predictions of the p197 repeat region. A) A single 175-residue repeat modeled alone, 30 superposed models: both programs on each of two registers of the repeat period, residues 414-588 and 579-753, with 5 models per AlphaFold 3 job and 10 per AlphaFold 2 job. One model is colored by pLDDT, with band boundaries at 50, 70 and 90 as keyed; the remaining 29 are shown in grey. Scale bar, 10 nm. B) Six tandem repeats, residues 414-1463, AlphaFold 2. Three of the five models are shown: the longest, the median and the shortest by end-to-end distance. All five appear in Figure S8C under the same numbering. Each model is centered in its own frame; the three frames share one scale, which differs from that of A, so the single scale bar applies to all three. Color identifies the repeat and not the confidence. The black line within each repeat is its fitted principal axis; the angles in C are measured between consecutive axes. Scale bar, 10 nm. C) Predicted aligned error against junction angle, for every model of the two- and six-repeat constructs, with the single-repeat models of A in the column marked “alone”. Filled symbols, the error between two consecutive repeats (65 junctions), plotted at the angle between the principal axes of the two repeats; 0° corresponds to a collinear continuation and 180° to a complete fold-back. Open symbols, the same quantity within a single repeat of the same models. Symbol shape indicates the construct, as keyed: circles, the 50 junctions of the six-repeat models; squares, the 15 junctions of the two-repeat models (Table S2). The column marked “alone” (diamonds) gives the within-repeat error of the 30 single-repeat models of A, which have no junction. Dashed line, maximum of the PAE scale, 31.73 Å.

In tandem, the repeats remain largely helical, but the models do not converge on the orientation of one repeat relative to the next. Figure 6B shows three of the five AlphaFold 2 six-repeat models, the longest, the median and the shortest by end-to-end distance (43.5, 14.6 and 4.8 nm), each repeat colored separately and carrying its fitted principal axis. Figure 6C plots the predicted aligned error, which measures how confidently one part of a model is placed relative to another, against the angle between consecutive axes (0°, a collinear continuation; 180°, a complete fold-back). Over the 65 junctions of the two- and six-repeat models the angle spans 3 to 178°, most junctions lying near one extreme or the other. The error between consecutive repeats is 26.1 to 31.4 Å, on a scale saturating at 31.73 Å, against 15.6 to 22.0 Å within a repeat of the same models and 12.1 to 15.2 Å in the 30 models of Figure 6A (full matrices, Figure S8A). Confidence also falls overall in the six-repeat models, to a mean pLDDT of 58 to 75 (Table S1), with the lowest values at the repeat boundaries (Figure S8B). All ten six-repeat models are shown in Figure S8C, D, with their geometry in Table S2. Given that the C- and N-termini of p197 are anchored on opposite sides of the exclusion zone and that the central repeat section of p197 makes up most of the protein, the filaments observed in our tomograms likely correspond to this central region. The measured EZF length (mean ∼417 nm) is approximately half of the ∼920 nm expected if the 35 repeats of 175 amino acids were arranged as a fully extended α-helix.

We conclude that p197 forms a flexible filament whose central body consists of an array of α-helical repeat units, flanked by unstructured terminal domains that anchor the (p)BB to the downstream TAC components at the mitochondrion.

## Discussion

Cryo-ET of *T. brucei* allowed us to image the cytoplasmic part of the TAC in its native context. We resolved the EZFs connecting the (p)BB to the OMM, identified the DMs, and measured the dimensions and orientations of these components across many cells. Two features stood out: a wide spread of (p)BB positions and angles relative to the OMM, and EZF lengths spanning ∼230 to 625 nm in zoid cells and ∼29 to 874 nm in NP40-treated cells. Together with prior evidence that p197 alone defines the (p)BB to OMM distance (Aeschlimann et al., 2022), these observations point to p197 as a flexible, length-adjustable connector and provide the basis for a structural model of how it can absorb the mechanical demands of BB movement.

Cryo-ET addressed two limitations of classical thin-section TEM. First, ultramicrotomy rarely captures the BB, the pBB, and a central section of the kDNA within the same cell, and when it does, the structures are mostly aligned along the section plane (Ogbadoyi et al., 2003). Cryo-ET on intact cells removed this orientation bias and revealed structural details that 2D imaging cannot capture. Second, it allowed us to image any cell in the sample without preselection at low magnification, avoiding the detection bias inherent to manually searching for regions of interest in fixed, sectioned samples. The overall organization we observed matched expectations from prior TEM studies of the TAC region (Ogbadoyi et al., 2003). Two differences in appearance reflect the absence of chemical fixation and heavy-metal staining: cell membranes appeared smoother than in chemically fixed and resin-embedded samples, and the kDNA was less electron-dense (Figure S4).

Within the TAC region, we identified the EZFs in approximately 30% of our tomograms. The number of visible filaments varied from a few to several dozen per cell. We attribute this variability primarily to technical limitations (low contrast due to fluctuations in cell thickness, ice quality, and the precision of tilt-series alignment) rather than to a true absence of filaments. In contrast, the ULFs were not visible in any of our tomograms. This likely reflects the similar mass density of the ULFs and the surrounding mitochondrial matrix, which is denser than the cytoplasm due to its high protein concentration (Kühlbrandt, 2015).

Our measured BB-to-OMM distance (145 ± 123 nm) is notably shorter than the ∼270 nm reported by Aeschlimann et al. (2022) using ultrastructure expansion microscopy (U-ExM), and shorter still than their measured length of p197 itself (∼200 nm). Part of this difference likely reflects the two techniques: U-ExM and cryo-ET involve different sample preparation and could introduce different measurement biases. More fundamentally, however, the two studies measure different quantities. Our BB-to-OMM distance is a straight-line gap between the (p)BB and the OMM, whereas our measured EZF length (417.5 nm) is a contour length traced along the full path of a wavy, curving filament. These are not directly comparable: a filament can readily span a distance far shorter than its own contour length if it is not held taut. Indeed, the fact that a ∼417 nm filament typically bridges a gap of only ∼145 nm is itself informative, indicating that the EZFs are not stretched tight between the (p)BB and the OMM but instead run a slack or curved path between their two anchor points, consistent with the flexible, length-variable behavior we propose for p197.

The pBB sat closer to the OMM than the mature BB on average, with the BB position showing greater spread (63 ± 81 nm vs 145 ± 123 nm; Figure 3D). This pattern is consistent with the mature BB subjected to flagellar beating while the pBB, which lacks an attached flagellum, remains more constrained.

In the DM area, the OMM and IMM were separated by 8.2 ± 0.9 nm, compared to 9.9 ± 1.3 nm in adjacent regions of the mitochondrion (a 17% reduction; Figure 3C). This narrowing is consistent with the inner and outer membranes being physically tethered in the TAC region by the inner-membrane-spanning TAC protein p166 and the outer-membrane component TAC60 (Stettler et al., 2024).

Beyond the continuous perturbation from flagellar beating discussed above (Figure 7A), the BB experiences two additional movements during the cell cycle. The first is the rotation of the new BB/pBB pair around the old BB during kinetoplast S-phase (Figure 7B), which repositions the new BB/pBB pair posterior to the parental BB (Lacomble et al., 2010; Vaughan & Gull, 2015). This rotation occurs at late stages of pBB outgrowth, when the transition zone is fully formed, (Lacomble et al., 2010; Vaughan & Gull, 2015) and a new set of pBBs has nucleated next to the old flagellum and the freshly matured BB (Vaughan & Gull, 2015). Our tomograms show that p197 assembles at the new BB before these events, with filaments already present at the maturing BB before nucleation of the new set of pBBs (Figures 1 and 2). The TAC connection therefore exists during BB rotation, and any flexibility of the filaments must accommodate this movement. The second cell cycle event is the central function of the TAC: physical segregation of the replicated kDNA networks. After outgrowth of the new axoneme (Figure 7C), and completion of kDNA replication, the two BB/pBB pairs separate (Robinson & Gull, 1991), and because the TAC links each BB to a daughter kDNA, this separation directly drives kDNA segregation into the daughter. All three sources of movement (flagellar beating, rotation, and separation) impose mechanical demands on the TAC, on the attached membranes, and on the kDNA itself.

**Figure 7:**
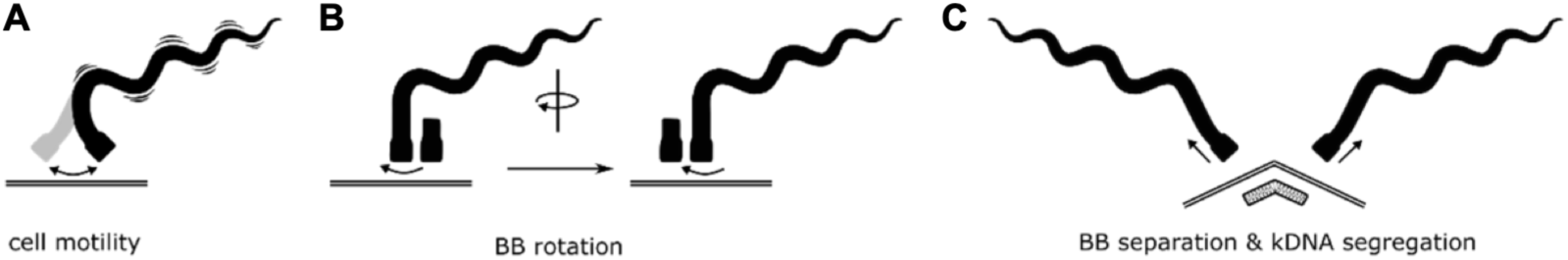
Impact of cell motility and cell cycle dynamics on the TAC. A) through C), shown are different types of BB/pBB movement relative to the OMM. A) Being the base of a motile flagellum, the mature BB is subject to slight but constant tumbling movements. B) The freshly matured BB has been shown to rotate around its maternal BB to relocate to a position posterior of the maternal BB. C) kDNA segregation depends on the separation of the BBs prior to cytokinesis. A stable connection between the BBs and the kDNA is essential for kDNA segregation.

How does p197 accommodate the mechanical demands placed on the TAC? Our EZF measurements show that traced filament contour lengths vary. Within a single tomogram, the five filaments differed by 158 nm on average and by up to 282 nm, and across the zoid dataset they ranged from 229.6 to 624.5 nm. Because those five filaments belong to the same basal body at the same moment, the variation cannot reflect differences between cells or between cell cycle stages. The repeat section, some 35 nearly identical α-helical units in tandem, is a natural substrate for it: a length change distributed over many equivalent units requires only a small change in each. What that change is at the molecular level is not resolved by our data.

The predictions constrain the secondary structure of the repeat but not the arrangement of consecutive repeats (Figure 6), and this limit follows from the sequence data available to them. The multiple sequence alignment of the p197 repeat region contains 29 sequences and is never more than 19 deep at any position (Figure S8E). p197 has few homologues in the sequence databases, and the orthologues found in other kinetoplastids lack the repeat section altogether (Aeschlimann et al., 2022). No covariation signal is therefore available from which the packing of one helix against another could be inferred. The models fall into two regimes: where the repeats are built as extended helices the next is placed antiparallel to it, in all seventeen such junctions at 165 to 178°, and where they collapse the angle between them takes any value. Neither is asserted with confidence, the error between consecutive repeats staying near the top of its scale in both, so the range of angles records the absence of a determined geometry rather than a range of conformations. The absence of a converged inter-repeat geometry in the models is thus a property of the alignment rather than evidence for flexibility of the filament, and the flexibility we propose rests on the EZF length measurements.

Functional support for our model in *T. brucei* comes from loss-of-function data: a truncated version of p197 lacking the predicted α-helical region fails to maintain the kDNA, even though the truncated protein still localizes correctly and binds both the BB and the OMM (Aeschlimann et al., 2022). This separation of localization from function suggests that the helical region is not required for assembly into the TAC but is essential for its mechanical role. Consistent with this, replacing the repeat domain with the corresponding α-helical region of the *T. cruzi* ortholog, which has no repeats, does not restore function (Aeschlimann et al., 2022).

Several limitations remain. First, direct structural validation of our model awaits high-resolution determination of p197. Our attempts to purify recombinant fragments of the central repeat domain did not yield material suitable for cryo-EM or crystallography, leaving the precise architecture of the repeat unit undetermined. Second, we do not resolve how the repeat region changes length. Single-molecule force spectroscopy on isolated p197 fragments would provide the most direct test, measuring the force-extension behavior of the repeat array directly. Third, the ULFs were not resolved in our tomograms, and the molecular nature of the connection between the kDNA and the inner mitochondrial membrane therefore remains structurally uncharacterized. Mitochondrial permeabilization to reduce matrix density may make these filaments accessible to cryo-ET in future studies.

In summary, cryo-ET of the *T. brucei* TAC reveals an architecture in which p197, the largest TAC protein in *T. brucei*, anchors the kDNA to the BB, while accommodating the mechanical demands of basal body movement, which drives kDNA segregation during the cell cycle. This combination of flexibility and stable anchorage may represent a general design strategy for cellular structures that link rigid organelles across mechanically dynamic cellular contexts.

## Materials and Methods

### *T. brucei* cell culture conditions

Procyclic form 29-13 asynchronous *T. brucei* cells were cultured in semi-defined medium-79 (SDM-79) supplemented with 10 % FCS, 15 µg/ml geneticin and 25 µg/ml hygromycin at 27° (Wirtz et al. 1999). The cell line is part of the established collection of the Institute of Cell Biology, University of Bern, Bern, Switzerland.

From the 29-13 cell line, the two RNAi cell lines TbCentrin4 (Tb927.7.3410) RNAi and TbCentrin4 p197 (Tb927.10.15750) double RNAi were generated. Depending on the cell line, 5 µg/ml phleomycin and/or 10 µg/ml blasticidin was added to the media (see below). For RNAi induction, 1 μg/ml tetracycline was used.

### Cloning of RNAi construct

To clone the TbCentrin4 RNAi construct, we inserted base pairs 19-438 of the ORF (TREU927) into the pFC-4 vector. The sequence was obtained by PCR: FWD-primer: 5’ – GTAAAAGCTTGGATCCGAACAGATCCGTGAAGCG – 3’; REV-primer: 5’ GTAATCTAGACTCGAGCATCTGCATCATGACGCTC – 3’; template DNA: NYsm DNA isolate. Using the restriction sites (HindIII, BamHI, XbaI and XhoI) introduced by the primers, the target sequence was inserted twice (in opposite direction) to encode for a hairpin dsRNA. The pFC-4 plasmid contains blasticidin resistance. To clone the p197 RNAi construct, a fragment of the p197 ORF was inserted into the pZJM vector by PCR amplification and inserted twice (in opposite orientation) to encode a hairpin dsRNA. The pZJM plasmid contains phleomycin resistance.

### *T. brucei* transfections

To obtain the TbCentrin4 RNAi and TbCentrin4 p197 double RNAi cell lines, we transfected cells with the constructs described above. We integrated the constructs by homologous recombination. 10^8^ PCF cells or TbCentrin4 RNAi cells were used. The transfection mixtures consisted of 10 μg of linearized plasmid in 110 μl transfection buffer (90 mM sodium phosphate, pH 7.3), 5 mM KCl, 0.15 mM CaCl_2_, 50 mM HEPES, pH 7.3) (Burkard et al., 2007). Cells were pelleted at 2’500 rcf for 8 min, mixed carefully with the transfection mixture and electroporated with the Amaxa Nucleofector (program X-014) (Schumann Burkard et al., 2011). After transfection, the cells were recovered in antibiotic free SDM-79 for 20 h. After the recovery, we added the respective antibiotics to select for integration of the transfected constructs. To select for TbCentrin4 RNAi, we used 10 µg/ml blasticidin. For p197 RNAi, 5 µg/ml phleomycin were used.

### Immunofluorescence analysis

10^6^ cells were pelleted for 3 min at 1800 rcf. Cells were washed with 1 ml PBS, resuspended in 20 μl PBS, and spread on a glass slide. After settling, cells were fixed for 4 min with 4% paraformaldehyde in PBS. Following fixation, they were permeabilized for 5 min with 0.2% Triton-X 100. The sample was then blocked with 4% bovine serum albumin in PBS for 30 min. Cells were incubated with primary and secondary antibodies (diluted in blocking solution) for 45-60 min as follows: rat YL1/2 antibody detecting tyrosinated tubulin, which is found in the BB (Kilmartin et al., 1982)(a kind gift of Keith Gull) 1:10’000, monoclonal mouse anti-TAC102 antibody (Trikin et al., 2016) 1:5’000, Alexa Fluor® 488 goat anti-rat IgG (H+L)(Nanoprobes/FluoroNanogold) 1:1000, Alexa Fluor® 647 goat anti-mouse IgG (H+L) (Life technologies) 1:1000. After each antibody incubation step, cells were washed 3 x with 0.1 % Tween-20 (in PBS). A final wash with PBS was performed before mounting the cells in ProLong® Gold Antifade Mounting medium with DAPI (4’,6-diamidine-2-phenylindole) (Invitrogen).

### Fixed section transmission electron microscopy

Trypanosomes were grown as described above, harvested and centrifuged at 2500 rcf for 5 min. The pellets were fixed with 2.5 % glutaraldehyde in 0.15 M HEPES pH 7.41 at 4° C for at least 24 h. They were then washed with 0.15 M HEPES three times for 5 min, post-fixed with 1% OsO4 in 0.1 M sodium cacodylate buffer at 4° C for 1 h, washed with 0.05 M maleate-NaOH buffer three times for 5 min. Subsequently the cells were dehydrated in 70, 80, and 96 % ethanol for 15 min each, at room temperature. Then the cells were immersed in 100% ethanol three times for 10 min, in acetone two times for 10 min, and finally in acetone-epon (1:1) overnight, at room temperature. Cells were then embedded in pure epon and left to harden at 60° C for 5 days. Ultrathin sections (70−80 nm) were produced with an ultramicrotome UC6 (Leica Microsystems). The sections, mounted on 200 mesh copper grids, were stained with uranyless and lead citrate with an ultrostainer (Leica Microsystems). Sections were then examined with a transmission electron microscope (Tecnai Spirit, 80 kV) equipped with an Olympus-SIS Veleta CCD camera.

### Sample preparation for cryo-electron tomography of detergent treated cells

PCF cells were harvested during exponential growth at a concentration of 5×10^6^ cells/ml. For the subsequent permeabilization experiments, 10^7^ cells were used. Permeabilization was done as described previously (Sasse & Gull, 1988). In short, Ethylenediaminetetraacetic acid (EDTA) was added to a final concentration of 0.5 mM followed by centrifugation at 2000 rcf for five minutes. Subsequently the supernatant was removed, and the cells were resuspended in extraction buffer (10 mM NaH_2_PO_4_, 150 mM NaCl, 1mM MgCl_2_, 0.5% NP40 or 0.5% Triton X-100, pH 7.2) to a concentration of 5×10^8^ cells/ml and incubated on ice for seven minutes (NP40) or 10 minutes (Triton X-100). The cells were centrifuged at 3000 rcf for three minutes and resuspended in PBS to a final concentration of 5 · 10^8^ cells/ml. Glow-discharged cryo-EM grids (Cu, R2/1, Quantifoil) were prepared by applying three microliters of the permeabilized cells (Vitrobot, three seconds blotting time, 15 seconds incubation, 100% humidity).

### Sample preparation for cryo-electron tomography of zoids

TbCentrin4 RNAi or TbCentrin4 p197 double RNAi PCF cells were RNAi induced for 65 h. The cells were then harvested by centrifugation at 2500 rcf, for 8 min. They were then resuspended in serum-free SDM-79 to a concentration of 2 · 10^7^ cells/ml, and treated with a series of centrifugation steps: 26 rcf for 2 min, 68 rcf for 3 min, 210 rcf for 5 min. Then, the upper 35 % of the sample were collected for further centrifugation: 210 rcf for 3 min, 340 rcf for 5 min. We then collected the upper 55 % of this fraction, before centrifuging again, at 345 rcf for 3 min. At this point we collected the upper 60 % of the sample, and pelleted all cells remaining in this faction, by centrifugation at 1800 rcf for 7 min. The cells were washed in PBS, and then resuspended to a final concentration of 4 · 10^7^ cells/ml. 10 nm gold beads (Aurion, OD520 = 2.0) were added 1:10 v:v. 3 μl of this sample were placed on a glow-discharged EM grid (lacey carbon films on Cu 200 mesh, Quantifoil Micro Tools), blotted from the back side for 3 s, and plunged into liquid ethane at a temperature of ≤ -170° C. The grids were stored in liquid nitrogen, until further use.

### Sample preparation with focused ion beam milling

Cells were blotted on EM grids (R2/1 holey carbon 200 films on Cu 200 mesh, Quantifoil Micro Tools). The grids were first glow-discharged and 4 µl of trypanosome wild-type culture (2-4 · 10^7^ cells/ml) were placed on the grid. Using a Vitrobot Mark 4 (Thermo Fisher Scientific), cells were plunge-frozen in a liquid ethane-propane mixture after blotting for 12 s at blotting force -4. Afterwards, grids were clipped into autogrid support rings with a cut-out that allows access to the ion beam at a low angle (Thermo Fisher Scientific) and stored in liquid nitrogen until being used for FIB milling.

Cryo-FIB milling was performed as described previously (Schaffer et al., 2017) with an Aquilos 2 dual-beam FIBSEM instrument (Thermo Fisher Scientific). In the FIB/SEM chamber, grids were coated with a layer of organometallic platinum using a gas injection system to protect the sample surface. Micro-expansion joints (relief cuts) were milled to prevent lamella from bending. A gallium ion beam was used for milling (Wolff et al., 2019). Owing to the small size of individual trypanosome cells, small clusters of cells were milled at a low angle (14-15°) to form short lamellae of 70-200 nm thickness. After milling, grids were transferred into a Titan Krios transmission electron microscope (Thermo Fisher Scientific), operating at 300 kV with a post-column energy filter (GIF Quantum LS, Gatan), and a direct detector camera (K2, Gatan). A total electron dose of 60-80e^-^/Å^2^ was distributed over a tilt range of -58° to +58°, with an increment of 2°. SerialEM (V4.0) was used for data collection.

### Data acquisition and processing of cryo-electron tomograms

The vitrified samples were screened at 200 kV to assess the ice quality. Tilt-series were acquired on Titan Krios transmission electron microscope with an operational voltage of 300 kV; a Volta phase plate; a Falcon4 (Thermo Fisher Scientific) direct electron detector and the Thermo Fisher Scientific Tomography 5 software (V5.22). A dose-symmetric scheme was applied in 2° increments ranging from -58° to +58° with a total dose of 120 e^-^ /Å^2^. The acquired tilt-series were processed using AreTomo3 (V2.1.5). Segmentation was performed using IMOD (V5.1.0), with the resulting data imported into Blender (v4.2 LTS) using MicroscopyNodes (V2.1.1) for rendering and visualization. All figures were made using Adobe Illustrator (v2026).

Tilt-series were acquired on Titan Krios transmission electron microscopes (Thermo Fisher Scientific) operating at 300 kV. The microscopes used were equipped with a post-column energy filter, K2 (Gatan), K3 (Gatan), or Falcon4 (Thermo Fisher Scientific) direct electron detector, and in some cases, volta phase plate was used to enhance contrast at low defocus. Using SerialEM, tomograms were acquired as dose-symmetric tilt-series in 2° increments, ranging from -58° to +58°. A total electron dose of 120 e^-^/Å^2^ was applied.

After acquisition, tilt-series of cryo-FIB milled trypanosomes were preprocessed using the TOMOMAN pipeline (v.0.6.9) (Khavnekar et al., 2024) and custom MATLAB scripts. Raw frames of all tilt-series (including cryo-FIB milled trypanosomes as well as TbCentrin4 zoids and tdzoids) were then aligned using MotionCor2 (v1.3.2 or v.1.4.7) 2017). Then, each cryo-electron tomogram was reconstructed in IMOD (v4.12.0) (Mastronarde & Held, 2017; S. Q. Zheng et al., 2017) analyzed visually, and tomograms were selected for further processing. This processing included deep learning-based denoising using cryo-CARE (v0.1.1) (Buchholz et al., 2018) and in a few cases, manual segmentation of structures of interest, using IMOD. Tomograms from cryo-FIB milled trypanosomes were further processed with IsoNet (v1), to compensate for missing wedge artifacts. Different IMOD versions were used because of the timely difference in tomogram acquisition (Liu et al., 2022). In a separate pipeline, tilt series were aligned and reconstructed with AreTomo3 (S. Zheng et al., 2022), and the resulting tomograms were denoised with CryoSamba (Costa-Filho et al., 2025), which substantially improved the visibility of subset of tomograms.

Distance measurements were conducted with the “measure” function of the IMOD drawing tools. Filament length was measured by placing contours on the relevant structures and reading out of all contour lengths of the respective IMOD model. Filament diameter was measured from manually selected stretches of well-resolved filaments. The local filament diameter was then measured from the intensity curve along a line drawn orthogonally to the filament direction. The effective diameter was defined as the width of the intensity drop at half its depth. Intermembrane distance was measured using the FIJI macro (https://forum.image.sc/t/imagej-macro-to-measure-distance-between-two-lines-edges/42019). This macro takes two manually segmented lines as an input to return the distance between the two lines at 15 positions along them. The measured values correspond to the distance between centers of the respective membrane.

Angle measurements were performed using a custom-made python script. Upon entry of the coordinates of four points and a plane (defined by three points) in 3D space, the script calculates the normal vector of the plane, and the directional vectors of points one and two, and of points three and four, respectively. Based on the normal vector of the plane, and the directional vectors, it then further calculates the angle between each directional vector and the plane, as well as the angle between the two directional vectors. To apply this to our dataset, we defined the mitochondrial membranes in the TAC area as the plane, and the BB and pBB as directional vectors. We manually collected the respective points needed to define 3D orientation of the three objects of interest in the reconstructed tomograms.

(p)BB distance to the mitochondrial membranes, angle measurements and filament length were measured on denoised tomograms acquired without volta phase plate, at defocus values ranging from -2 to -10 μm. EZF diameter was measured on non-filtered, unbinned (pixel size: 0.434 nm) tomograms acquired with volta phase plate, at defocus values of no more than -2.5 μm.

### Structure prediction of the p197 repeat region

The 175-residue repeat unit of p197 (TriTrypDB Tb927.10.15750) was modeled alone and in tandem with two programs. AlphaFold 2 was run through ColabFold 1.6.2 (Mirdita et al., 2022) using the alphafold2_multimer_v3 preset, three recycles, no templates, and multiple sequence alignments generated by MMseqs2 against UniRef and the environmental databases. AlphaFold 3 (Abramson et al., 2024) was run on the AlphaFold Server at default settings, which returns five models per job. ColabFold was run with two seeds for the one- and two-repeat constructs, giving ten models each, and with one seed for the six-repeat construct. The ten runs gave 70 models in total, all of which are reported (Table S1).

Constructs were cut from the repeat region of a 2224-residue sequence comprising the N-terminal region, six repeats, and the C-terminal region. Three single-repeat constructs sample three registers of the 175-residue period: residues 414-588, 579-753 and 492-666, cut at positions 1, 166 and 79 of the period respectively, the last of these falling in the middle of the long helix. The two- and six-repeat constructs comprised residues 579-928 and 414-1463.

Geometry was measured on the Cα trace. A residue was counted as helical when it belonged to a Cα(i)-Cα(i+4) span of 6.2 ± 1.2 Å. Repeats were taken as successive 175-residue blocks counted from the first residue of the construct, which corresponds to the repeat motif ERVEVATWDSEQQ. Two quantities were measured for each repeat: the length of its longest continuous helix, allowing breaks of up to 6 residues, and its span, the Cα5-Cα165 distance, which is 24 nm for a fully extended helix. A principal axis was fitted to each repeat by singular value decomposition over residues 5-165. The junction angle is the angle between the axis vectors of consecutive repeats taken N to C, 0° corresponding to a collinear continuation and 180° to a complete fold-back. Predicted aligned error was averaged over the corresponding block of the matrix and symmetrized, over the same residues. Table S1 averages over every pair of different repeats, while Figure 6C and Table S2 report consecutive pairs only. The scale of both programs saturates at 31.73 Å.

Geometric measurement of basal body and mitochondrial membrane orientation

For each tomogram, seven reference points were manually placed in the reconstructed volume: two points on the principal axis of the BB (P₁, P₂), two on the principal axis of the pBB (P₃, P₄), and three non-collinear points on the OMM in the TAC area (P₅, P₆, P₇).

The BB and pBB axis vectors were defined as v_BB = P₂ − P₁ and v_pBB = P₄ − P₃. The unit normal to the OMM plane was computed from the cross product (P₆ − P₅) × (P₇ − P₅), normalized to unit length and oriented according to the right-hand rule.

For each axis, the signed angle θ between the axis and the OMM plane was computed as

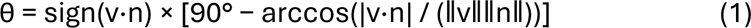

yielding values in [−90°, +90°]. By convention, θ > 0 indicates that the axis points away from the mitochondrial matrix and θ = 0 indicates an axis lying in the membrane plane. Because the order of the three OMM points is arbitrary, the initial direction of n is ambiguous. We exploited the biological constraint that both BB and pBB axes typically point from inside the cell toward the OMM (θ > 0). When both signed angles were negative, n was inverted and the angles recomputed. The angle between the BB and pBB axes was computed as the unsigned arccosine of the normalized dot product v_BB · v_pBB, yielding values in [0°, 180°].

Dot products were clipped to [−1, 1] before applying across to avoid numerical out-of-domain errors. All computations were performed in Python 3.10 using NumPy.

### Statistical analysis

Geometric annotations were collected on 73 tomograms. For each of the three angles (BB-to-OMM, pBB-to-OMM, BB-pBB), we report count, mean, median, and standard deviation. Distributions were visualized as histograms. The BB-OMM and pBB-OMM inclination distributions were compared using a Wilcoxon signed-rank test. Pairwise associations between the three angles were assessed using Pearson product-moment correlation, with two-sided p-values obtained from the asymptotic t-distribution implementation in SciPy. A significance threshold of α = 0.05 was used; no correction for multiple comparisons was applied, given the small and pre-specified number of tested associations.

Intermembrane distances in the TAC area and non-TAC region (Figure 3C), and BB and pBB distances to the OMM (Figure 3D), were each measured within the same cells, and compared using a two-sided paired t-test.

All statistical analyses were performed in Python 3.10 using pandas (data handling), NumPy, scipy.stats (pearson, wilcoxon), matplotlib (visualization), and R (v4.6.1).

For filament lengths, the tomogram was used as the unit of analysis, because filaments traced within one tomogram are not independent. The mean filament length was computed for each tomogram, and each dataset is summarized by the mean and sample standard deviation of its tomogram means, with a 95 % confidence interval from the t distribution on n − 1 degrees of freedom; the range quoted for each dataset is that of the individual filament measurements. The zoid dataset comprises 110 filaments in 22 tomograms (five traced per tomogram) and the NP40 dataset 430 filaments in 12 tomograms (11 to 88 traced per tomogram). The two were analyzed separately throughout and compared on tomogram means, using Welch’s unequal-variance t test and the Mann-Whitney U test for location and Levene’s test for dispersion.

## Acknowledgments

This work was mainly funded by project grants to Torsten Ochsenreiter from Uniscientia and the Swiss National Science Foundation (grant number 179454) and by a project grant to Benoît Zuber from the Swiss National Science Foundation (grant number 179520). We acknowledge Diamond for access and support of the cryo-EM facilities at the UK national Bio-Imaging Centre (eBIC), proposals BI26915 and NT21004, funded by the Wellcome Trust, MRC and BBSRC. Equipment supported by the Microscopy Imaging Center (MIC) of the University of Bern, the eBIC of the Diamond Light Source, the BioEM facility of the University of Basel and ScopeM of the ETH Zürich was used during this study. Contributions from Takashi Ishikawa were funded by the grant number 310030_192644 of the Swiss National Science Foundation.

## Supplementary Materials, Tables and Figures

### Quantification of p197 repeat number by Southern blot

The central repeat domain of p197, the largest protein in *T. brucei*, has been difficult to resolve by short-read genome assembly due to the high sequence identity between individual 525-nucleotide repeats. The original genome assembly reported approximately 3.5 central repeats (Berriman et al., 2005), whereas a long-read cDNA assembly later estimated approximately 26 repeats (Naguleswaran et al., 2021). To determine the repeat number experimentally, we generated an engineered allele, p197*, by introducing a HindIII site followed by a 924-nucleotide recoded region upstream of the start codon. This recoded sequence, which does not cross-hybridize with the endogenous repeats, served as the specific binding site for the Southern blot probe. NheI sites occur naturally once per repeat within the central repeat domain, and a native XhoI site lies downstream of the repeat stretch, allowing repeat number to be assessed by partial digestion.

Genomic DNA was digested with HindIII and XhoI to release the complete p197* fragment. To generate a ladder of fragments corresponding to discrete repeat numbers, samples were subjected to either a complete NheI digest (2 h) or partial NheI digests (10 min) across a dilution series (1/45 to 1/1000), in addition to the HindIII/XhoI digest. Digested DNA was separated by agarose gel electrophoresis alongside a DNA ladder and analyzed by Southern blot using the probe against the recoded region.

Fragment sizes were determined by constructing a calibration curve from the migration distances of the DNA ladder bands (n = 3 lanes; mean ± SD, with a 90% confidence band fit to the data). The migration distance of the HindIII/XhoI-digested p197* fragment, measured across three independent experiments, was interpolated against this calibration curve to estimate its size, with the corresponding 68% confidence range reported. The number of central repeats was then inferred by subtracting the length of the recoded region and flanking non-repetitive sequences from the estimated fragment size and dividing the remainder by 525 bp per repeat, yielding an estimated repeat number and ORF length for the p197* locus with associated 68% confidence intervals.

**Table S1:** Summary of the AlphaFold predictions of the p197 repeat region.

| Program | Construct | n | Mean pLDDT | pTM | Longest helix [aa] | Span [nm] | PAE within [Å] | PAE between [Å] |
| --- | --- | --- | --- | --- | --- | --- | --- | --- |
| AF3 | 1 repeat, motif phase, 414-588 | 5 | 89.5 | 0.46 | 158 | 22.8 to 23.2 | 12.1 | n/a |
| AF2 | 1 repeat, motif phase, 414-588 | 10 | 83.6 to 88.1 | 0.42 to 0.44 | 158 to 160 | 23.7 to 24.0 | 13.3 to 15.2 | n/a |
| AF3 | 1 repeat,<br>helix phase,<br>579-753 | 5 | 86.2 to<br>86.6 | 0.44 to<br>0.45 | 160 | 23.5 to<br>23.7 | 13.0 to<br>13.4 | n/a |
| AF2 | 1 repeat,<br>helix phase,<br>579-753 | 10 | 83.3 to<br>90.3 | 0.43 to<br>0.45 | 160 | 23.8 to<br>24.0 | 12.5 to<br>15.0 | n/a |
| AF3 | 1 repeat,<br>mid-helix<br>phase, 492-666 | 5 | 72.8 to<br>76.0 | 0.35 to<br>0.36 | 129 to 160 | 13.6 to<br>23.7 | 19.9 to<br>20.2 | n/a |
| AF2 | 1 repeat,<br>mid-helix<br>phase, 492-666 | 10 | 60.6 to<br>71.6 | 0.37 to<br>0.41 | 160 | 1.6 to<br>10.5 | 17.1 to<br>19.6 | n/a |
| AF3 | 2 repeats,<br>helix phase,<br>579-928 | 5 | 68.8 to<br>69.6 | 0.28 | 133 to 160 | 15.0 to<br>23.7 | 15.6 to<br>16.9 | 30.1 to<br>30.8 |
| AF2 | 2 repeats,<br>helix phase,<br>579-928 | 10 | 45.5 to<br>74.2 | 0.26 to<br>0.31 | 107 to 160 | 6.8 to<br>22.9 | 16.4 to<br>20.0 | 26.1 to<br>29.6 |
| AF3 | 6 repeats,<br>414-1463 | 5 | 69.4 to<br>74.8 | 0.19 to<br>0.21 | 151 to 160 | 2.7 to<br>23.5 | 18.5 to<br>20.9 | 29.9 to<br>30.9 |
| AF2 | 6 repeats,<br>414-1463 | 5 | 58.4 to<br>66.1 | 0.18 to<br>0.20 | 93 to 160 | 4.5 to<br>13.5 | 19.6 to<br>21.7 | 30.4 to<br>31.0 |

AlphaFold 2 is ColabFold 1.6.2 with the alphafold2_multimer_v3 preset; AlphaFold 3 is the AlphaFold Server at default settings. Residue numbering refers to the 2224-residue construct. n is the number of models, and ranges span all models of a run, none excluded. Longest helix and span were measured over residues 5-165 of each repeat, so a repeat forming one fully extended helix scores 160 aa and 24 nm. PAE within is the mean over a single repeat. PAE between is the mean over every pair of different repeats, consecutive or not; Figure 6C and Table S2 report the same quantity for consecutive pairs only, which for the six-repeat models spans a wider range, 28.6 to 31.4 Å against 29.9 to 31.0 Å here. The alignment contained 29 sequences throughout and was never more than 19 deep at any position.

The three single-repeat constructs sample three registers of the 175-residue period: 414-588 begins at the repeat motif, 579-753 one residue after the inter-repeat loop, and 492-666 in the middle of the long helix. Figure 6A superposes the models of the first two. In the third, both termini fall within a helix, which accounts for the lower confidence of both programs and for the reduced span of the AlphaFold 2 models.

**Table S2:** Geometry of the individual two- and six-repeat models.

| Run | Model | End to end<br>[nm] | Longest helix<br>[aa] | Repeat span<br>[nm] | Junction angles<br>[°] |
| --- | --- | --- | --- | --- | --- |
| AF2, 6 rep | (i) | 43.5 | 93 to 159 | 5.5 to 12.7 | 9, 24, 53, 74, 74 |
| AF2, 6 rep | (ii) | 37.2 | 97 to 160 | 4.5 to 7.7 | 19, 22, 55, 39, 27 |
| AF2, 6 rep | (iii) | 14.6 | 149 to 157 | 7.2 to 11.5 | 37, 88, 156, 69,<br>129 |
| AF2, 6 rep | (iv) | 9.7 | 120 to 156 | 7.6 to 12.3 | 150, 155, 132,<br>44, 114 |
| AF2, 6 rep | (v) | 4.8 | 95 to 156 | 11.9 to 13.5 | 164, 178, 162,<br>177, 172 |
| AF3, 6 rep | (i) | 4.8 | 160 | 22.2 to 23.5 | 178, 166, 165,<br>176, 168 |
| AF3, 6 rep | (ii) | 4.2 | 152 to 160 | 22.4 to 23.3 | 171, 172, 178,<br>171, 169 |
| AF3, 6 rep | (iii) | 1.2 | 160 | 8.5 to 9.0 | 174, 177, 173,<br>174, 173 |
| AF3, 6 rep | (iv) | 3.8 | 151 to 158 | 5.3 to 6.6 | 5, 6, 3, 5, 4 |
| AF3, 6 rep | (v) | 0.8 | 158 | 2.7 to 2.8 | 9, 9, 9, 8, 9 |
| AF2, 2 rep | rank_003 | 18.5 | 113 to 160 | 9.2 to 11.0 | 23 |
| AF2, 2 rep | rank_006 | 13.8 | 112 to 160 | 6.8 to 9.5 | 9 |
| AF2, 2 rep | rank_005 | 12.2 | 107 to 113 | 6.9 to 8.3 | 10 |
| AF2, 2 rep | rank_002 | 10.2 | 160 | 10.8 to 19.1 | 169 |
| AF2, 2 rep | rank_008 | 9.0 | 158 to 160 | 11.3 to 17.6 | 176 |
| AF2, 2 rep | rank_010 | 7.6 | 112 to 160 | 12.5 to 16.3 | 172 |
| AF2, 2 rep | rank_001 | 7.5 | 160 | 19.9 to 22.9 | 165 |
| AF2, 2 rep | rank_004 | 5.2 | 160 | 12.9 to 14.0 | 171 |
| AF2, 2 rep | rank_007 | 5.1 | 160 | 11.1 to 14.3 | 167 |
| AF2, 2 rep | rank_009 | 3.2 | 141 to 160 | 16.9 to 18.1 | 178 |
| AF3, 2 rep | model_4 | 12.0 | 160 | 15.2 to 23.4 | 172 |
| AF3, 2 rep | model_1 | 11.8 | 160 | 15.3 to 23.5 | 170 |
| AF3, 2 rep | model_2 | 11.6 | 133 to 160 | 15.0 to 23.3 | 175 |
| AF3, 2 rep | model_3 | 11.3 | 137 to 160 | 15.4 to 23.4 | 171 |
| AF3, 2 rep | model_0 | 5.7 | 155 to 160 | 22.1 to 23.7 | 176 |

Longest helix and repeat span are given as the range over the repeats of each model, both measured over residues 5-165 of each repeat; a repeat forming one fully extended helix scores 160 aa and 24 nm. The six-repeat models carry the numbering used in the figures. The AlphaFold models are ordered by end-to-end distance, and Figure 6B shows (i), (iii) and (v); the AlphaFold 3 models are ordered by repeat span, their end-to-end distances all lying between 0.8 and 4.8 nm. The two-repeat models appear in no figure and are identified by their model number.

Figure S1: Additional cryo-electron tomograms of NP40-treated cells. A-F) Z-slices through cryo-electron tomograms of the TAC region from six NP40-treated cells, additional to the example shown in Figure 1. Visible are the p197 filaments (black arrowhead), the basal body (BB), the pro-basal body (pBB), the outer and inner mitochondrial membrane (OMM, IMM), and the scale bar: 100 nm.

Figure S2: Comparison of sample preparation methods to visualize TAC and p197 in PCF cells. A) Whole cell (untreated). B) An example of a zoid cell. C-D) Example of NP40 treated cells. E) An example of a tomogram of a cell treated with Triton X-100. F) FIB-milled whole wild type cell. Visible structures, depending on the preparation, include p197 filaments, the basal body (BB), the pro-basal body (pBB), the flagellar pocket (FP), the outer and inner mitochondrial membrane (OMM, IMM, MM). Scale bars: 100 nm. Images are averages of ten tomogram slices.

Figure S3: Sample preparation workflow and characterization of zoid and tdzoid cells A) Workflow for cryo-ET on cryo-FIB milled whole cells. (i) WT PCF cells are mounted onto carbon coated copper grids and (ii) plunge-frozen in liquid ethane. (iii) Lamellae of 70-200 nm are milled with a cryo-FIB scanning electron microscope. (iv) Tilt-series of low dose micrographs are acquired through the lamellae on a cryo-TEM, then (v) reconstructed and (vi) denoised (see Methods for details). The micrograph in (i) shows an overlay of phase contrast with the DNA stain DAPI; scale bar: 5 µm. B) Workflow for cryo-ET on enriched zoid and tdzoid populations. *T. brucei* cells carrying the TbCentrin4 RNAi construct or TbCentrin4/p197 double RNAi construct are (i) induced for 65 h, then (ii) enriched for zoids by differential centrifugation, (iii) plunge-frozen, and (iv-vi) acquired and processed as in A. Micrographs in (i) and (ii) show an overlay of phase contrast with the DNA stain DAPI; scale bars: 5 µm. C) Karyotype distributions of WT *T. brucei* (black), TbCentrin4 RNAi cells induced for 65 h (zoids, dark grey) and the corresponding enriched fraction (zoids enriched, light grey). n ≈ 200 cells per condition. D) Karyotype distributions of WT *T. brucei* (black), TbCentrin4/p197 double RNAi cells induced for 65 h (tdzoids, dark grey), and the corresponding enriched fraction (tdzoids enriched, light grey). n ≈ 150 cells per condition.

Figure S4: Characterization of zoid and tdzoid cells by immunofluorescence and conventional TEM. A) Immunofluorescence on WT *T. brucei* cells (top), zoid cells (middle), and tdzoid cells (bottom). The cells were immunostained with the BB marker YL1/2 (red), TAC marker TAC102 (green), and DAPI, which stains nuclear and kinetoplast DNA. Phase contrast (PH) shows the overall cell morphology. Zoid cells retain the BB, the TAC, and the kDNA (single DAPI spot, no nucleus), while tdzoid cells retain the BB but lose the TAC and the kDNA. Scale bar: 5 µm. B) TEM analysis of the TAC region in chemically fixed, WT resin-embedded cell (top) and a zoid cell (bottom). Scale bar: 250 nm.

Figure S5: Cryo-electron tomograms of tdzoid cells lacking p197. Three z-slices through a representative tomogram of a tdzoid cell (TbCentrin4/p197 double RNAi), showing the region between the BB and the mitochondrion. Arrows indicate the proximal face of the BB, where EZFs emanate in zoid cells but are absent here. Scale bar: 200 nm. Bottom row: enlarged views of the boxed regions, showing the area proximal to the BB. Scale bar: 100 nm.

Figure S6: The EZFs accommodate a wide range of basal body orientations relative to the mitochondrial membranes in zoid PCF cells. Top row: three representative tomographic slices showing the BB, pBB, and mitochondrial membranes (MM) in different orientations. Scale bar: 100 nm. Bottom row: schematic representations of the corresponding tomograms, highlighting the positions of these structures.

Figure S7: p197 contains about ∼35 repeats and is the largest protein in *T. brucei* A) Schematic of the p197 locus across two genome assemblies. The original assembly (Berriman et al., 2005) reported ∼3.5 central repeats, whereas a long-read gDNA assembly (Naguleswaran et al., 2021) reported ∼26 repeats. B) Strategy for quantifying repeat number by Southern blot on the engineered allele p197*, using a recoded probe-binding region upstream of the repeats and NheI sites within each repeat. C) Calibration curve for fragment size estimation from the gel in (E), with the interpolated size of the HindIII/XhoI-digested p197* fragment (red marker, 68% CI). D) Estimated repeat number and ORF length of the p197* locus, with 68% confidence intervals. E) Representative Southern blot of the p197* locus. Lane 1: HindIII/XhoI digest. Lane 2: additional complete NheI digest, showing the probe-proximal fragment (0 repeats). Lanes 3–6: additional partial NheI digests at four dilutions, resolving discrete fragments corresponding to increasing repeat numbers.

Figure S8: Confidence of the AlphaFold predictions of the p197 repeat region. A) Predicted aligned error of a six-repeat model, shown in full for one model of each program: the highest-ranked AlphaFold 2 model, which is (ii) in C, and the first AlphaFold 3 model, which is (i) in D. Lines mark the repeat boundaries and the axes are numbered by repeat; dark indicates high confidence. Only the diagonal blocks, corresponding to the repeats themselves, are informative. B) pLDDT along the chain for one repeat modeled alone and for the six-repeat construct, AlphaFold 3 in both cases and over the same residues. Vertical lines mark the repeat boundaries. C) All five AlphaFold 2 models of the six-repeat construct, drawn each centered in its own frame, and ordered by end-to-end distance; Figure 6B shows (i), (iii) and (v). D) All five AlphaFold 3 models of the same construct, drawn at the same scale as C and ordered by repeat span, their end-to-end distances all lying between 0.8 and 4.8 nm. Per-model geometry is given in Table S2. Scale bars, 10 nm. E) Number of sequences aligned at each position of the six-repeat construct. The search returned 29 sequences, and no position is more than 19 deep.

