## Supplementary Figures and Tables for "Cryo-electron tomography sheds light on the largest protein in the *Trypanosoma brucei* tripartite attachment complex"

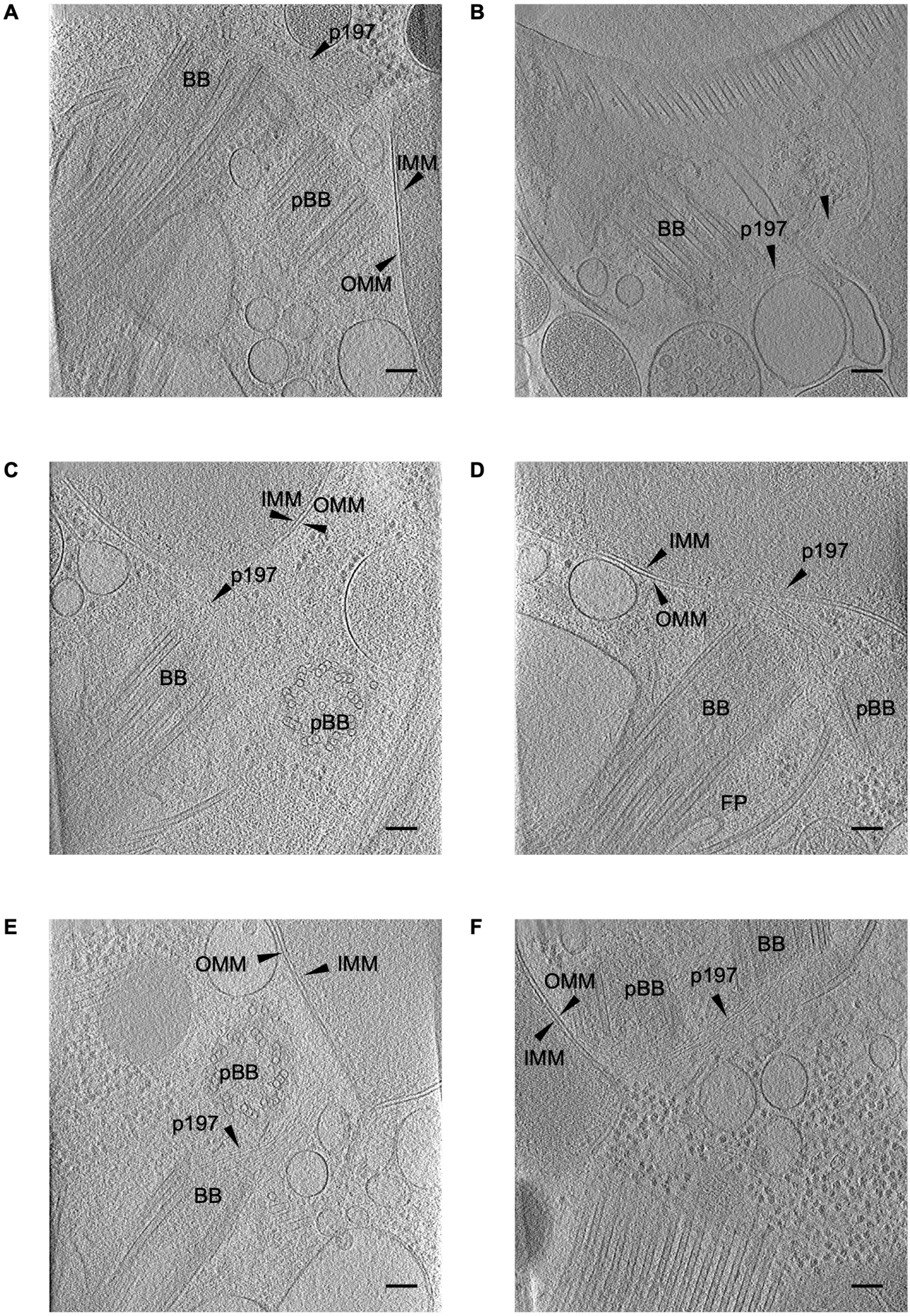
A
p197
BB
IMM
pBB
OMM
B
BB
p197
C
IMM
OMM
p197
BB
pBB
D
IMM
p197
OMM
BB
pBB
FP
E
OMM
IMM
pBB
p197
BB
F
BB
OMM
pBB
p197
IMM

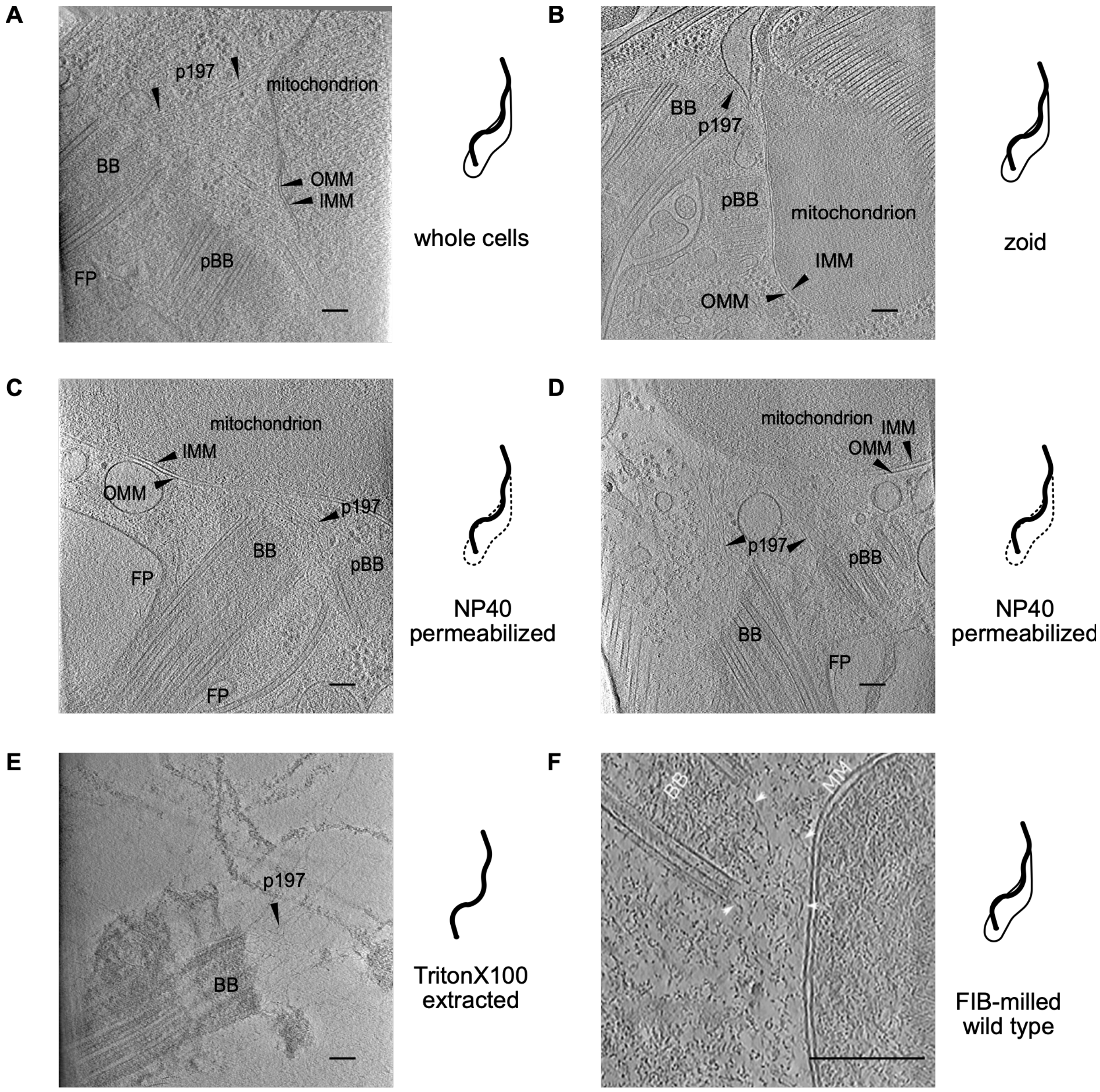
A
p197
mitochondrion
BB
OMM
IMM
FP
pBB
whole cells
B
BB
p197
pBB
mitochondrion
IMM
OMM
zoid
C
mitochondrion
IMM
OMM
p197
BB
pBB
FP
FP
NP40
permeabilized
D
mitochondrion
IMM
OMM
p197
pBB
BB
FP
NP40
permeabilized
E
p197
BB
TritonX100
extracted
F
BB
MM
FIB-milled
wild type

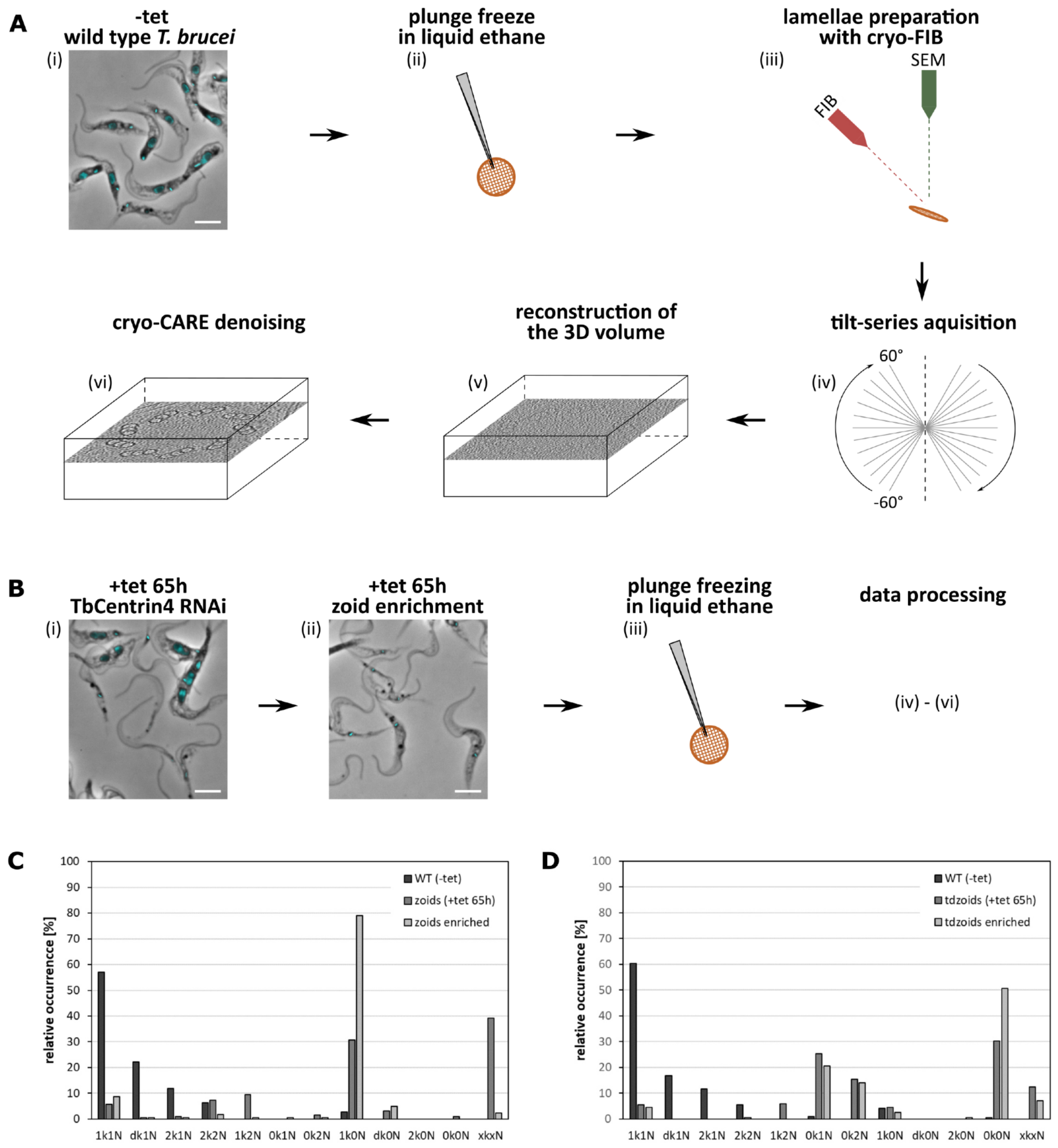
A
-tet
wild type T. brucei
(i)
plunge freeze
in liquid ethane
(ii)
lamellae preparation
with cryo-FIB
(iii)
SEM
FIB
tilt-series aquisition
(iv)
60°
-60°
reconstruction of
the 3D volume
(v)
cryo-CARE denoising
(vi)
B
+tet 65h
TbCentrin4 RNAi
(i)
+tet 65h
zoid enrichment
(ii)
plunge freezing
in liquid ethane
(iii)
data processing
(iv) - (vi)
C
relative occurrence [%]
WT (-tet)
zoids (+tet 65h)
zoids enriched
1k1N dk1N 2k1N 2k2N 1k2N 0k1N 0k2N 1k0N dk0N 2k0N 0k0N xkxN
D
relative occurrence [%]
WT (-tet)
tdzoids (+tet 65h)
tdzoids enriched
1k1N dk1N 2k1N 2k2N 1k2N 0k1N 0k2N 1k0N dk0N 2k0N 0k0N xkxN

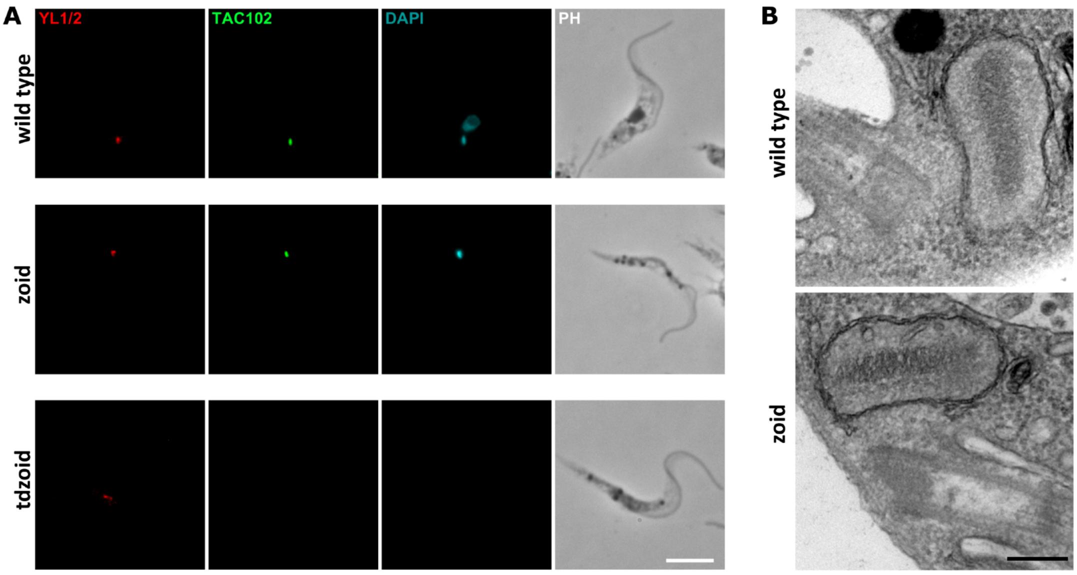
A
YL1/2
TAC102
DAPI
PH
wild type
zoid
tdzoid
B
wild type
zoid

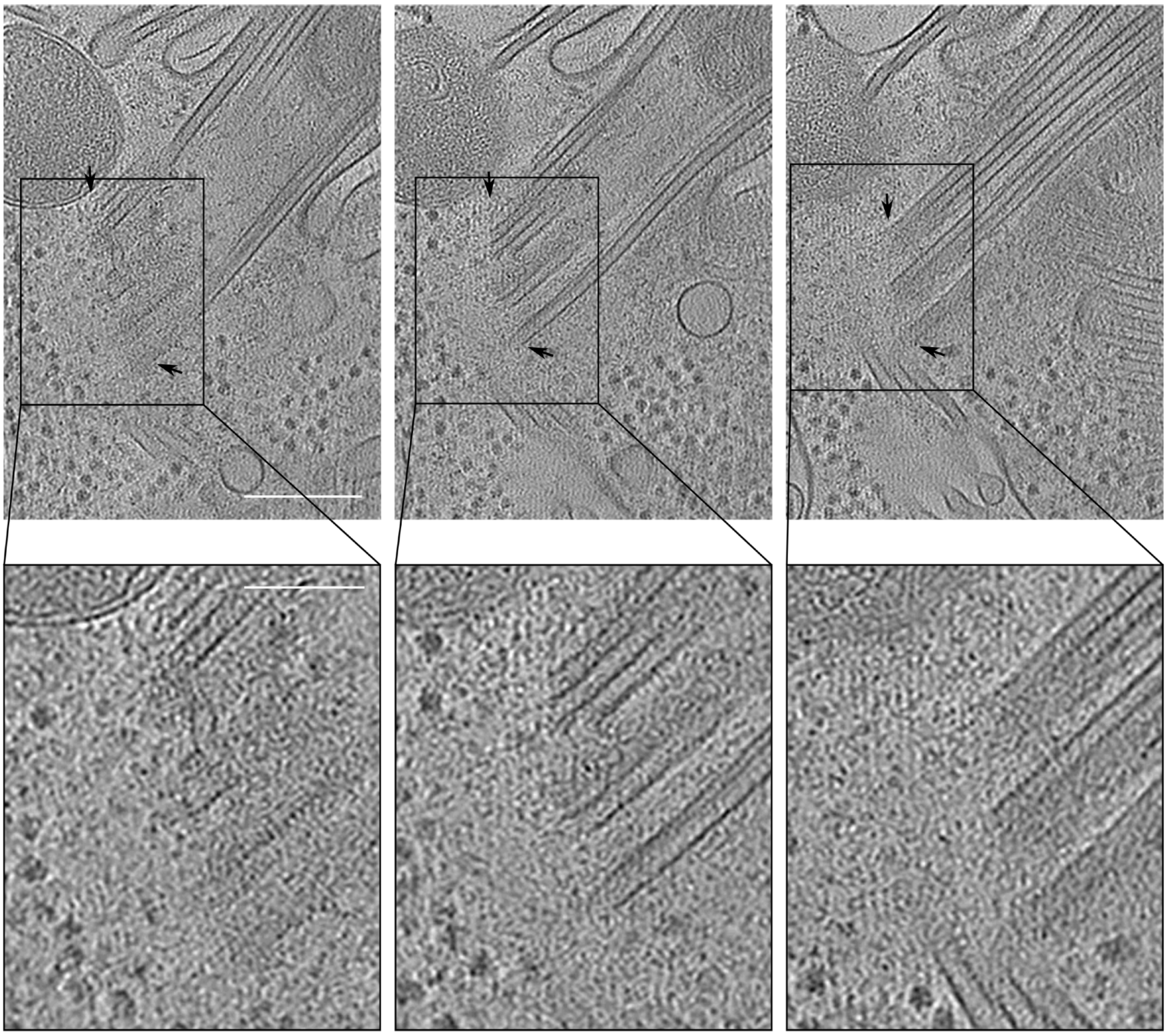

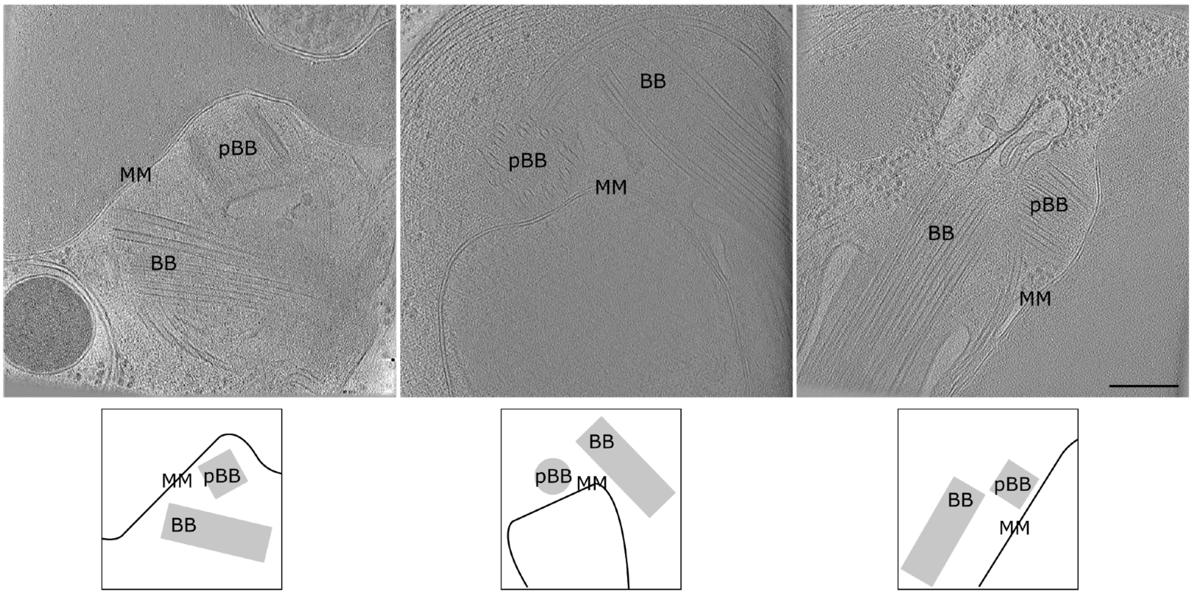
pBB
MM
BB
BB
pBB
MM
pBB
BB
MM
MM
pBB
BB
BB
pBB
MM
BB
pBB
MM

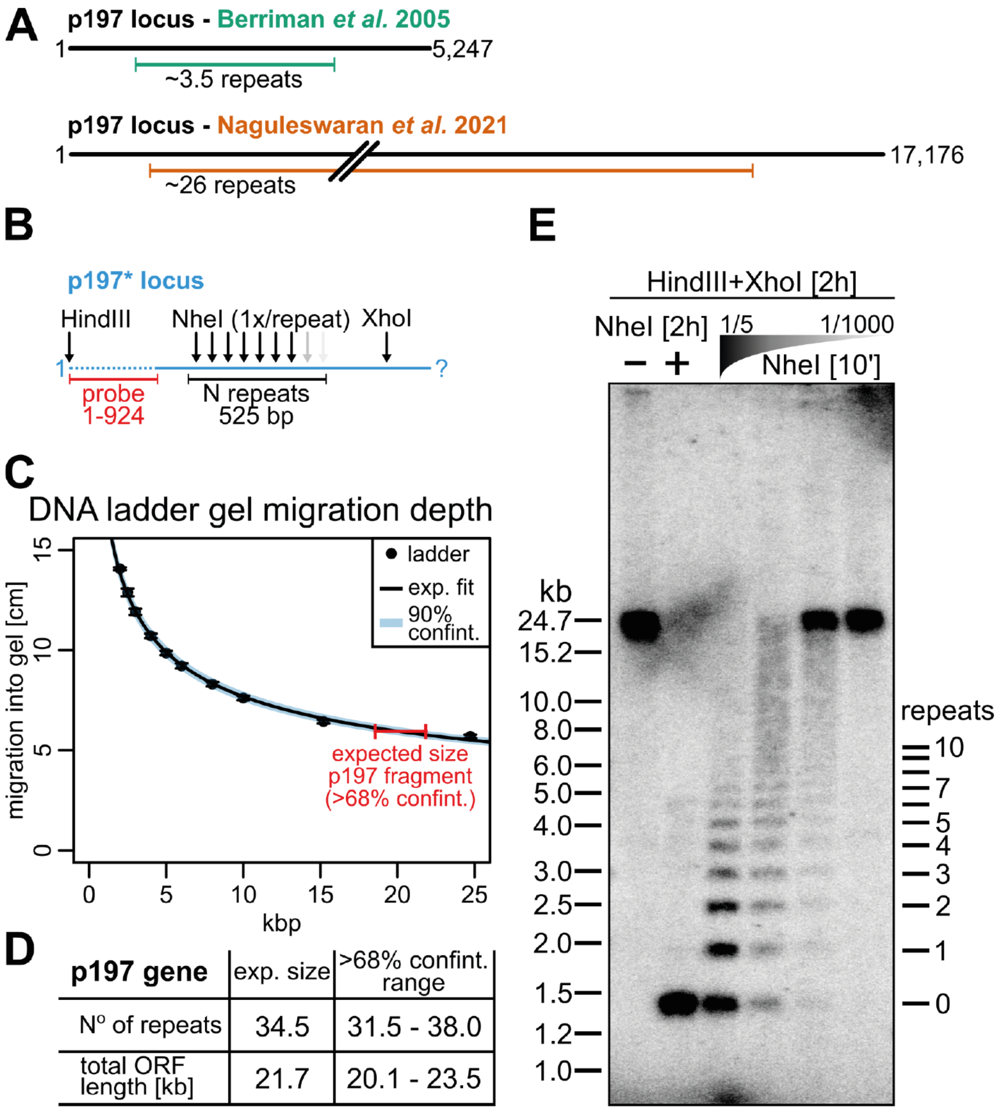


D

| p197 gene | exp. size | >68% confint. range |
|---|---|---|
| N° of repeats | 34.5 | 31.5 - 38.0 |
| total ORF length [kb] | 21.7 | 20.1 - 23.5 |

(A) p197 contains a repetitive central domain encoded by 525 nucleotide long nearly identical sequences. The first complete genome sequencing (Berriman et al 2005) resulted in an assembly with ~3.5 repeats (top) while a long-read cDNA sequencing study (Naguleswaran et al. 2021) produced an assembly with ~26 repeats.
(B) To estimate the number of repeats in the p197 locus we introduced a HindIII site directly upstream of the start codon followed by a 924 nucleotide long recoded sequence allowing the specific detection of this engineered allele. The p197 locus contains naturally NheI sites (1x per repeat) and and XhoI site downstream of the repeat stretch.
(C) Genomic DNA of the engineered cell line (B) was digested with HindIII and XhoI over night and DNA fragments were separated on 0.6% agarose gels. p197 fragments were selectively probed with a radioactively-marked oligomer complementary to the recoded region of the engineered p197 locus. The migration distance of (n=3) marker lanes and (n=3) p197 fragments were fitted to estimate the size of the HindIII/XhoI digested p197 locus fragment.
(D) Expected repeat number and 68% confidence range as well as expected p197 locus length with 68% confidence range calculated from the fitted data in (C).
(E) Example of a Southern blot probed for the engineered p197 locus. Left 1 (from left): Fragment produced by HindIII/XhoI digest. Lanes 2-6: Fragments produced by HindIII/XhoI complete digests together with a complete digest by NheI (lane 2) or concentration-dependant incomplete digests by NheI (lanes 3-6). Discrete signals in lanes 3-6 correspond to p197 locus fragments with increasing numbers of repeats.

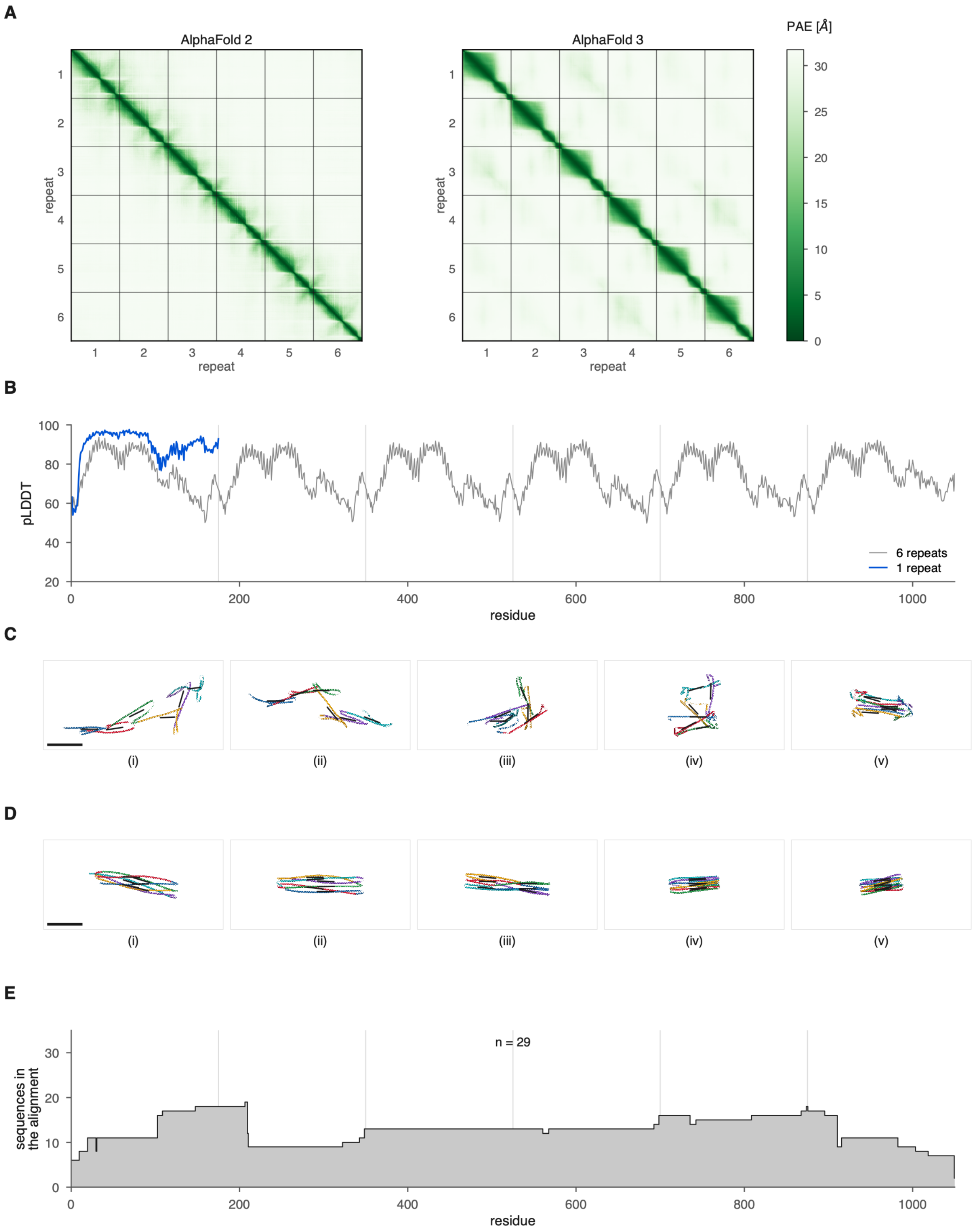
A
AlphaFold 2
AlphaFold 3
PAE [Å]
30
25
20
15
10
5
0
repeat
1
2
3
4
5
6
B
pLDDT
100
80
60
40
20
0
200
400
600
800
1000
residue
6 repeats
1 repeat
C
(i)
(ii)
(iii)
(iv)
(v)
D
(i)
(ii)
(iii)
(iv)
(v)
E
sequences in the alignment
n = 29
30
20
10
0
0
200
400
600
800
1000
residue
